# *Aluminum Resistance Transcription Factor 1 (ART1)* contributes to natural variation in rice Al resistance

**DOI:** 10.1101/137281

**Authors:** Juan David Arbelaez, Lyza G. Maron, Timothy O. Jobe, Miguel A. Piñeros, Adam N. Famoso, Ana Rita Rebelo, Namrata Singh, Qiyue Ma, Zhangjun Fei, Leon V. Kochian, Susan R. McCouch

**Affiliations:** School of Integrative Plant Science, Plant Breeding and Genetics Section, Tower Rd, Cornell University, Ithaca, NY 14853; Robert W. Holley Center for Agriculture and Health, USDA-ARS, Tower Rd, Cornell University, Ithaca, NY 14853; Boyce Thompson Institute, Tower Rd, Cornell University, Ithaca, NY 14853

**Keywords:** rice, aluminum, abiotic stress, acid soils, QTL mapping, transcriptional regulation

## Abstract

Transcription factors (TFs) mediate stress resistance indirectly via physiological mechanisms driven by the genes they regulate. When studying TF-mediated stress resistance, it is important to understand how TFs interact with different genetic backgrounds. Here, we fine-mapped the aluminum (Al) resistance QTL *Alt12.1* to a 44 Kb region containing six gene models. Among them is *ART1*, which encodes a C2H2-type zinc finger TF required for Al resistance in rice. The parents of the mapping population, Al-resistant Azucena (*tropical japonica*) and Al-sensitive IR64 (*indica*), showed similar *ART1* expression levels but extensive sequence polymorphism within the *ART1* coding region. Using reciprocal near-isogenic lines (NILs) in the Azucena and IR64 genetic backgrounds, we examined how allele-swapping *Alt12.1* would affect plant responses to Al. Analysis of global transcriptional responses to Al stress in roots of the NILs alongside their recurrent parents demonstrated that the ART1 from Al-resistant Azucena led to greater changes in gene expression in response to Al when compared to the ART1 from IR64 in both genetic backgrounds. The presence of the ART1 allele from the opposite parent affected the expression of several genes not previously implicated in rice Al tolerance. We highlight examples where putatively functional variation in *cis*-regulatory regions of ART1-regulated genes interacts with ART1 to determine gene expression in response to Al. This ART1-promoter interaction may be associated with transgressive variation for Al resistance in the Azucena × IR64 population. These results illustrate how ART1 interacts with the genetic background to contribute to quantitative phenotypic variation in rice Al resistance.

## Introduction

Plants achieve abiotic stress resistance through diverse mechanisms that evolved through both natural and artificial selection. The characterization of quantitative trait loci (QTL) and their underlying genes has led to insights about the genetic architecture and the biological mechanisms of resistance to several abiotic stresses. Two major classes of genes predominate as determinants of stress resistance: membrane transporters and transcription factors (TFs) (Mickelbart et al., 2015). The reprogramming of gene expression through transcriptional regulation, mediated by TFs, is a finely orchestrated and tightly regulated process, and is one of the hallmarks of plant response to stress (Vaahtera and Brosché, 2011). To achieve specificity in the transcriptional response, a TF must activate only target genes involved in adaptation to a particular stress or combination of stresses (Vaahtera and Brosché, 2011). TFs are activated through post-translational modification, nuclear import and/or may be transcriptionally activated by other TFs. Genetic variation in the TF loci, or in any element along the regulatory chain, can contribute to phenotypic variation in plant stress resistance. Understanding the quantitative nature of TF-mediated stress resistance in plants is essential when predicting the impact of introducing alleles into a new genetic background in the context of plant breeding and crop improvement.

Aluminum toxicity severely limits plant growth on acidic soils (pH <5). Under these conditions, the rhizotoxic Al species Al^3+^ is solubilized, inhibiting root growth and function (Kochian et al., 2015), and leaving plants more vulnerable to drought and mineral nutrient deficiencies. Approximately 30% of the earth’s land area consists of highly acid soils, and as much as 50% of all potentially arable lands are acidic (von Uexküll and Mutert, 1995). As vast areas of acid soils in the tropics and subtropics are critical food-producing regions, Al toxicity constitutes a food security threat exceeded only by drought among the abiotic limitations to crop production.

Rice (*Oryza sativa* L.) is the most Al-resistant species among the small grain cereals (Foy, 1988; Ma et al., 2002). A comparative cross-species study in hydroponics showed that rice is two-to six-fold more resistant than maize, wheat and sorghum (Famoso et al., 2010). This high level of resistance is likely achieved by the pyramiding of multiple mechanisms conferred by multiple genes, a hypothesis supported by results from both genome-wide association (GWAS) and QTL studies (Famoso et al., 2011). Mapping of Al resistance QTL in a rice recombinant inbred line (RIL) population derived from a cross between Al-resistant Azucena (*O. sativa* L., *tropical japonica*) and Al-sensitive IR64 (*O. sativa* L., *indica*) cultivars identified four genomic regions associated with Al resistance, on chromosomes 1, 2, 9, and 12 (Famoso et al., 2011; Spindel et al., 2013). The QTL *Alt12.1* on chromosome 12, which explains a large proportion (> 19%) of the variation in Al resistance, encompasses a genomic region that includes the gene *ALUMINUM RESISTANCE TRANSCRIPTION FACTOR 1* (*ART1*), encoding a C2H2-type zinc finger transcription factor (Yamaji et al., 2009). The *art1* mutant, producing a truncated version of the ART1 protein, is sensitive to Al stress.

A total of 32 genes were up-regulated in response to Al in the wild-type but were not up-regulated in the *art1* mutant (Yamaji et al., 2009). Some of these genes have been shown to play a role in rice Al resistance. These “ART1-regulated genes” have received this denomination because they are mis-regulated in the *art1* mutant; in other words, their expression is up-regulated by Al stress in the wild type, but not in the mutant (Yamaji et al., 2009). It is worth noting that direct binding of the ART1 protein to upstream regulatory regions has so far been experimentally demonstrated only for *STAR1* (Tsutsui et al., 2011) and *OsFRDL4* (Yokosho et al., 2016).

The genetic architecture of rice Al resistance is complex and quantitative in nature (Famoso et al., 2011). In the present study, we describe how natural variation in the *ART1* locus affects transcriptional responses to Al stress in two distinct genetic backgrounds of rice. We fine-mapped the Al resistance QTL *Alt12.1* to a small region surrounding *ART1* and used reciprocal near-isogenic lines (NILs) to examine the effect of two different *ART1* alleles in *japonica* (Al-resistant Azucena) and *indica* (Al-sensitive IR64) genetic backgrounds. Analysis of the global transcriptional response to Al stress in roots of the reciprocal NILs demonstrated that the *ART1* allele from Al-resistant Azucena induced greater changes in gene expression in response to Al stress when compared to the ART1 from IR64 in both genetic backgrounds. These changes were observed in a number of genes in addition to those previously reported as ART1-regulated. Moreover, our data suggest that the changes in gene expression pattern in response to the ART1 allele-swapping may be background-specific, suggesting that natural variation in *cis*-regulatory regions also plays a role in determining gene expression responses to Al. These results demonstrate the importance of genetic background in determining the phenotypic impact of a major Al resistance gene.

## RESULTS

### Development and evaluation of reciprocal near-isogenic lines (NILs) for the Al resistance QTL *Alt12.1*

We developed reciprocal near-isogenic lines (NILs) in the Azucena (Al-resistant) and IR64 (Al-sensitive) genetic backgrounds to examine how allele swapping at the *Alt12.1* QTL would affect plant responses to Al (Figure 1). The chromosomal region harboring *Alt12.1* from the Al-resistant parent Azucena was introduced into IR64 via backcrossing, and vice-versa (Methods S1). Phenotypically, the above-ground parts of AZU_[IR6412.1]_ and IR64_[AZU12.1]_ plants closely resemble their respective recurrent parents when grown under greenhouse conditions (Fig. 1A). The NIL AZU_[IR6412.1]_ contains a 3.92 Mb introgression from IR64 encompassing *Alt12.1* in the Azucena background, while the reciprocal NIL IR64_[AZU12.1]_ carries a 2.36 Mb Azucena introgression spanning the *Alt12.1* locus in the IR64 genetic background. Both NILs carry a single donor introgression across the target region, while over 99% of their genomes are identical to the recipient background (Fig. 1C; Table S2). When evaluated for Al resistance based on relative root growth (RRG) in hydroponics, the NIL AZU_[IR6412.1]_ was significantly (*p* < 0.05) less Al-resistant than its recurrent parent Azucena. Conversely, IR64_[AZU12.1]_, was significantly (*p* < 0.05) more Al-resistant than its recurrent parent, IR64 (Fig. 1B-D, Supplemental Table S2). These results validate the effect of the *Alt12.1* QTL on Al resistance in both genetic backgrounds.

**Figure 1:**
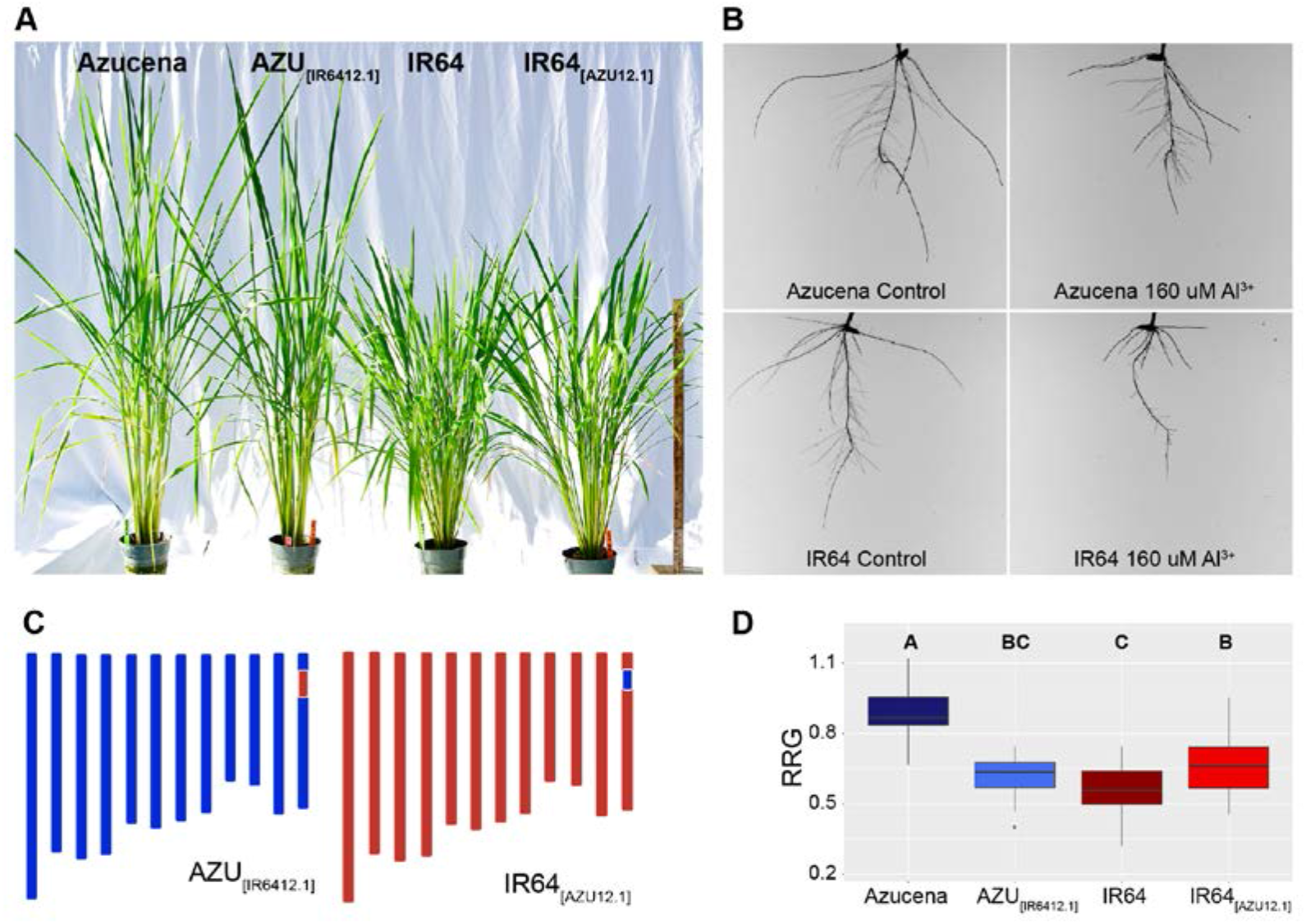
Near-isogenic lines (NILs) carrying reciprocal introgressions of the *Alt12.1* region validate the effect of the QTL on rice Al resistance. **(A)** Representative plants of the Al-resistant rice variety Azucena, of the Al-sensitive variety IR64, and of the reciprocal NILs near-isogenic lines AZU_[IR6412.1]_ and IR64_[AZU12.1]_ grown under greenhouse conditions. Photograph taken 110 days after sowing. **(B)** Parents of the recombinant inbred line (RIL) mapping population used to map Al resistance QTL (Famoso et al., 2011). Azucena (*tropical japonica*) is Al-resistant and IR64 (*indica*) is Al-sensitive. The photographs show roots of representative five day-old seedlings of Azucena and IR64 grown in hydroponics under control (0 μM Al^3+^ activity, left) and Al stress (160 μM Al^3+^ activity, right) condition for five days. **(C)** Genotypic makeup of the near-isogenic lines AZU_[IR6412.1]_ (left) and IR64_[AZU12.1]_ (right). In the schematic representation of the 12 chromosomes of rice, blue denotes Azucena background, and red denotes IR64 background. The reciprocal introgressions at the *Alt12.1* region on chromosome 12 are indicated by a blue rectangle in AZU_[IR6412.1]_ and by a red rectangle in IR64_[AZU12.1]_. **(D)** Al resistance phenotypes of Azucena, IR64, and the reciprocal NILs AZU_[IR6412.1]_ and IR64_[AZU12.1]_. Relative root growth (RRG) was calculated as the ratio between the total root growth (TRG) of seedlings (n = 18) grown under stress conditions (160 μM Al^3+^ activity) over TRG of seedlings (n = 18) grown under control conditions (0 μM Al^3+^ activity).

### Fine mapping of the *Alt12.1* QTL

The *Alt12.1* locus was fine-mapped using a substitution mapping approach (Methods S2) to a 0.2 cM region (44.7 Kb; chromosome 12: 3,578,363 - 3,623,299 bp), between markers *K_3.57* and *indel_3.62* (Figure 2A, Supplemental Figure S1, Supplemental Table S1). According to the rice MSUv.7 genome assembly (Kawahara et al., 2013), this region encompasses six gene models: *ART1* (LOC_Os12g07280), LOC_Os12g07290, LOC_Os12g07300, LOC_Os12g07310, LOC_Os12g07340, and LOC_Os12g07350.

**Figure 2:**
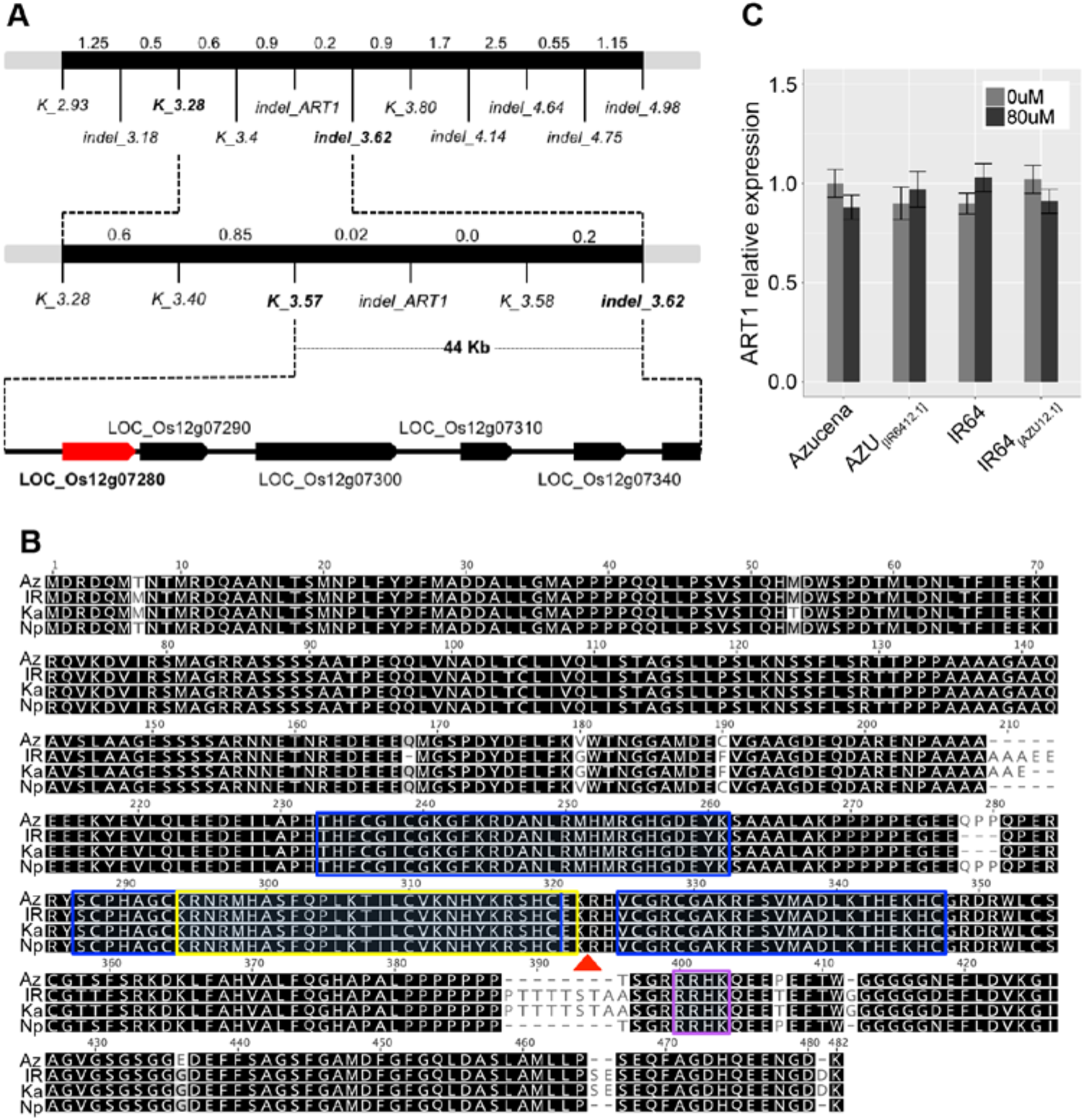
The rice Al resistance gene *AL RESISTANCE TRANSCRIPTION FACTOR 1* (*ART1*) localizes to the fine-mapped region of the Al resistance QTL *Alt12.1.* **(A)** Fine-mapping schema. The Al resistance QTL *Alt12.1* was fine-mapped using 4,224 F_3_ and 3,552 F_4_ plants from a cross between the Al-sensitive parent IR64 and an Al-resistant recombinant inbred line from the Azucena × IR64 mapping population (Supplemental Methods S2). The black bar represents a region of rice chromosome 12; molecular markers used at each step are indicated below the bar. Genetic distances are shown in pseudo cM above the black bar (1 cM ~ 200,000 bp). *ART1* (LOC_Os12g07280) is highlighted (red); other genes in the region are also represented (black arrows). **(B)** Alignment of the predicted ART1 amino acid sequences from Azucena, IR64, Kasalath, and Nipponbare. Predicted C2H2 domains (blue box) are shown; predicted monopartite (purple rectangle) and bipartite (yellow rectangle) nuclear localization domains are also shown. A red triangle under the alignment indicates the position of the 1 - bp deletion found in the *art1* mutant (Yamaji et al., 2009). **(C)** *ART1* relative expression in rice roots, measured using RT-qPCR. Azucena, IR64, and the reciprocal NILs AZU_[IR6412.1]_ and IR64_[AZU12.1]_ were grown in hydroponics and treated with Al (80 μM Al^3+^ activity) for 4 hours. Control: light-gray bar; Al stress, dark-gray bar. RNA was collected from six independent biological replicates. Error bars indicate a 95% confidence interval. All pairs of means were compared using a Tukey-Kramer HSD test and no significant differences were found (*p* < 0.05).

To assess the levels of structural variation between *japonica* and *indica* across the 44.7 Kb fine-mapped *Alt12.1* locus, we aligned sequences from the *japonica* rice reference genome, Nipponbare (which served as a proxy for Azucena) to a long-read, *de novo* assembly of the IR64 genome (Schatz et al., *in preparation*; http://schatzlab.cshl.edu/data/ir64/). A single IR64 scaffold was identified that covered > 36 Kb of the *Alt12.1* fine-mapped locus (44.7 Kb). No major rearrangements were detected in the region, and both gene order and gene content were preserved (Supplemental Figure S2). Based on previous reports showing the major role played by *ART1* in rice Al resistance, and the extreme Al sensitive phenotype of the *art1* mutant (Yamaji et al, 2009), we considered *ART1* to be the primary candidate gene underlying the Al resistance QTL *Alt12.1*.

### Allelic variation in *ART1*

To analyze the extent of allelic variation in the *ART1* gene, we examined both sequence and expression level polymorphisms. First, we sequenced the CDS in four rice varieties representing four major rice subpopulations: IR64 (*indica*), Azucena (*tropical japonica*), Kasalath (*aus*) and Nipponbare (*temperate japonica*) (Garris et al., 2005; Zhao et al., 2011). Azucena and Nipponbare, both Al-resistant *japonica* cultivars (Famoso et al., 2011), differ from each other by a single, non-synonymous amino acid substitution at position 436 of the ART1 protein (Fig. 2B). IR64 and Kasalath, both Al-sensitive cultivars, differ from each other by one non-synonymous substitution (position 53) and two indels (positions 168 and 210-211). The two *japonica* cultivars differ from the *indica* and *aus* cultivars by a total of 11 polymorphisms (4 aa substitutions and 7 indels). The most notable difference is a large C-terminal insertion (8 aa) that is present in IR64 and Kasalath (position 387-395) but absent in Azucena and Nipponbare. Between Azucena and IR64, the parents of the mapping population used in this study, we observed 4 aa substitutions and 7 indels (Fig. 2B).

Next we examined *ART1* expression levels in Azucena, IR64, and the reciprocal NILs AZU_[IR6412.1]_ and IR64_[AZU12.1]_. An RT-qPCR study in the presence and absence of Al found that *ART1* expression is not responsive to Al stress in the roots of Azucena, AZU_[IR6412.1]_, IR64, and IR64_[AZU12.1]_, and that there are no significant differences (*p* < 0.05) in *ART1* transcript accumulation in the roots of any of the four lines (Fig. 2C). These results suggest that the phenotypic variation associated with the *Alt12.1* QTL is likely due to genetic variation in the *ART1* coding region rather than expression level polymorphism.

### Functional analysis of the proteins encoded by different *ART1* alleles

In light of the polymorphisms observed in the *ART1* CDS, we were interested to determine whether the proteins encoded by the different *ART1* alleles differ in their nuclear targeting and/or DNA-binding affinity. *In-silico* analysis of the predicted ART1 proteins identified a bipartite nuclear localization signal upstream of the functional 1-bp frameshift mutation reported by Yamaji et al. (2009) in the *art1* mutant (Fig. 2B). This differed from the monopartite nuclear localization domain originally reported by Yamaji et al. (2009), which occurred downstream of the frameshift mutation. We used YFP fusions to determine the subcellular localization of the proteins encoded by the four natural *ART1* alleles as well as the aberrant protein expressed by the *art1* mutant. The *art1* mutant is characterized by a single-nucleotide deletion that disrupts the frame of translation at position 317, resulting in a longer protein (477 amino acids; the wild-type ART1 is 465 amino acids). The *art1* mutant version was generated via mutagenesis *in vitro* (see Materials & Methods). C-terminal YFP fusion proteins were expressed in tobacco leaves under control of a 35S promoter. A strong YFP signal was detected in the nucleus of cells of tobacco leaves expressing all four ART1 alleles as well as the *art1* mutant (Fig 3A). We conclude that the polymorphisms in the *ART1* coding region do not affect the subcellular localization of the protein.

**Figure 3:**
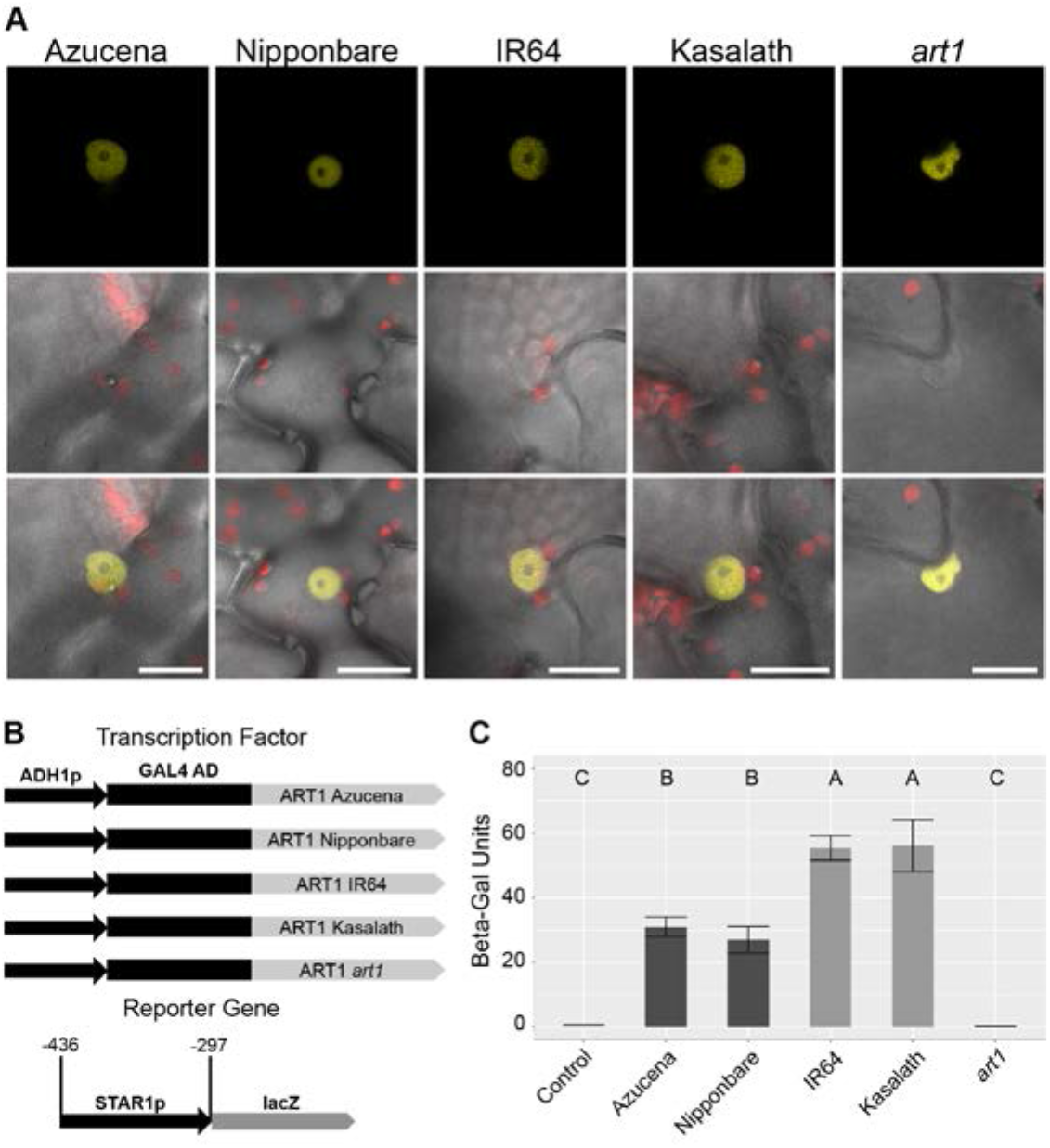
Proteins encoded by different *ART1* alleles localize to the nucleus, but differ in their DNA-binding affinity. **(A)** Subcellular protein localization of ART1 using C-terminal YFP gene fusions (ART1:YFP) transiently expressed in tobacco (*N. benthamiana*). Plasmids carrying the Azucena, Nipponbare, IR64, Kasalath, or *art1* mutant allele of ART1 were individually infiltrated into tobacco leaves. The top row shows the YFP fluorescence, while the middle row shows the superimposed chloroplast fluorescence and bright field channels. The third row shows merged images from all three channels (YFP fluorescence, chloroplast fluorescence, and bright field). Scale bar = 20 μm. **(B)** Schematic diagram of the constructs used in yeast one-hybrid assays. For the reporter construct, a fragment of the *STAR1* promoter was cloned in front of the *lacZ* gene and integrated into the yeast genome. Individual yeast colonies carrying the reporter construct were selected and transformed with one of five different transcription factor constructs. Each transcription factor construct contained the constitutive ADH1 promoter driving the expression of the GAL4 activation domain (GAL4 AD) fused to the *ART1* CDS from Azucena (*tropical japonica;* Al-resistant), Nipponbare (*temperate japonica*; Al-resistant), IR64 (*indica*; Al-sensitive), Kasalath (*aus* Al-sensitive), and the *art1* mutant allele. **(C)** Yeast one-hybrid assay comparing promoter activation by different *ART1* alleles. A 164-bp fragment of the *OsSTAR1* promoter was used. Constructs carrying an empty vector (control), or the Azucena, Nipponbare, IR64, Kasalath, or *art1* mutant allele were assayed for beta-galactosidase reporter gene activity (n = 10). Error bars indicate standard deviation. All pairs of means in **(C)** were compared using a Tukey-Kramer HSD test; lines not connected by the same letter are significantly different (*p* < 0.05).

Next we compared the DNA-binding ability of the proteins encoded by different *ART1* alleles in yeast one-hybrid assays (Fig. 3B). To evaluate the DNA-binding ability of the Nipponbare, Azucena, Kasalath, IR64 and *art1*-mutant alleles, we used stably transformed yeast lines carrying a fragment of the *OsSTAR1* promoter known to contain the ART1-binding site and previously used in yeast one-hybrid assays (Yamaji et al., 2009) to drive a beta-galactosidase reporter gene (Fig. 3B). When quantified via β-galactosidase assays, the *art1* mutant allele was unable to activate the reporter (Fig. 3C), in agreement with the fact that the 1-bp frameshift mutation in the *art1* mutant disrupts one of the predicted DNA-binding domains. The *japonica* alleles found in Nipponbare and Azucena, which differed by a single amino acid, showed comparable levels of reporter gene activation, indicating that they have similar DNA-binding affinity (Fig. 3C). The *aus* allele from Kasalath and the *indica* allele from IR64 activated the reporter gene at similar levels. On the other hand, the two *japonica* alleles (Azucena and Nipponbare) activated the reported gene at significantly lower levels (*p* < 0.05) when compared to the *indica/aus* alleles (IR64 and Kasalath).

### Whole transcriptome analysis in roots of Azucena, IR64 and the reciprocal NILs

We hypothesized that the polymorphisms also affect DNA-binding of the TF *in planta*, and its ability to activate downstream genes in response to Al stress. We examined this by performing transcriptome analysis in roots of Azucena, IR64 and the reciprocal NILs under Al stress using RNA-seq. Seedlings of Azucena, IR64 and the reciprocal NILs AZU_[IR6412.1]_ and IR64_[AZU12.1]_ were grown in hydroponics, then subjected to treatment with or without Al for 4 hours. Four independent biological replicates per genotype per treatment were collected and analyzed. A total of ~580M reads were generated across the 32 samples, with an average of ~18M reads per sample. After removing adaptors, low-quality sequences, and reads that align to ribosomal RNA, we uniquely mapped ~360M reads to the Nipponbare reference genome (MSUv.7), averaging 11M reads per sample (Supplemental Table S3). Raw read counts for each gene model were used to identify differentially expressed genes. The genotypic identity of all samples was confirmed by visualizing SNPs both within and outside the introgression regions (Suppl. Table S4). Differentially expressed genes were selected according to the following criteria: **a)** two-fold or greater change in transcript abundance; **b)** adjusted *p* value of ≤ 0.05; and **c)** minimum of eight counts in at least one sample.

Samples of each reciprocal NIL were analyzed alongside samples of their respective recurrent parents, to avoid spurious results due to genotypic differences across the *indica* and *japonica* genetic backgrounds. A principal component analysis (PCA) on normalized counts was performed to determine which variance components explain most of the global transcriptional variation. In the Azucena background (Azucena and NIL AZU_[IR6412.1]_), the first PC explains 44.5% of the variation and separates control from Al-treated samples (Supplemental Figure S3A). The second PC (23.7%) differentiates the two genotypes. In the IR64 background (IR64 and NIL IR64_[AZU12.1]_) the first PC explains 62% of the transcriptional variation, distinguishing the Al-treated IR64_[AZU12.1]_ samples from all others. The second PC (12.7%) separates the Al-treated IR64 samples from the control samples (Suppl. Fig. S3B). In both genetic backgrounds, the main source of transcriptional variation was Al treatment. However, in the IR64 background, the effect of Al treatment was more accentuated in the NIL IR64_[AZU12.1]_.

A total of 495 unique genes showed differential expression in response to Al across the four genotypes under study: 215 genes were up-regulated and 280 were down-regulated (Figure 4). The parental lines Azucena (*tropical japonica*, Al-resistant) and IR64 (*indica*, Al-sensitive) showed similar numbers of up-regulated genes (Azucena = 98; IR64 = 81), and half of them were shared across the two genotypes (n = 40; Fig. 4B). In contrast, Azucena showed a larger number of genes that were down-regulated in response to Al stress (n = 148) compared to IR64 (n = 35), and only 16 of them were shared.

**Figure 4:**
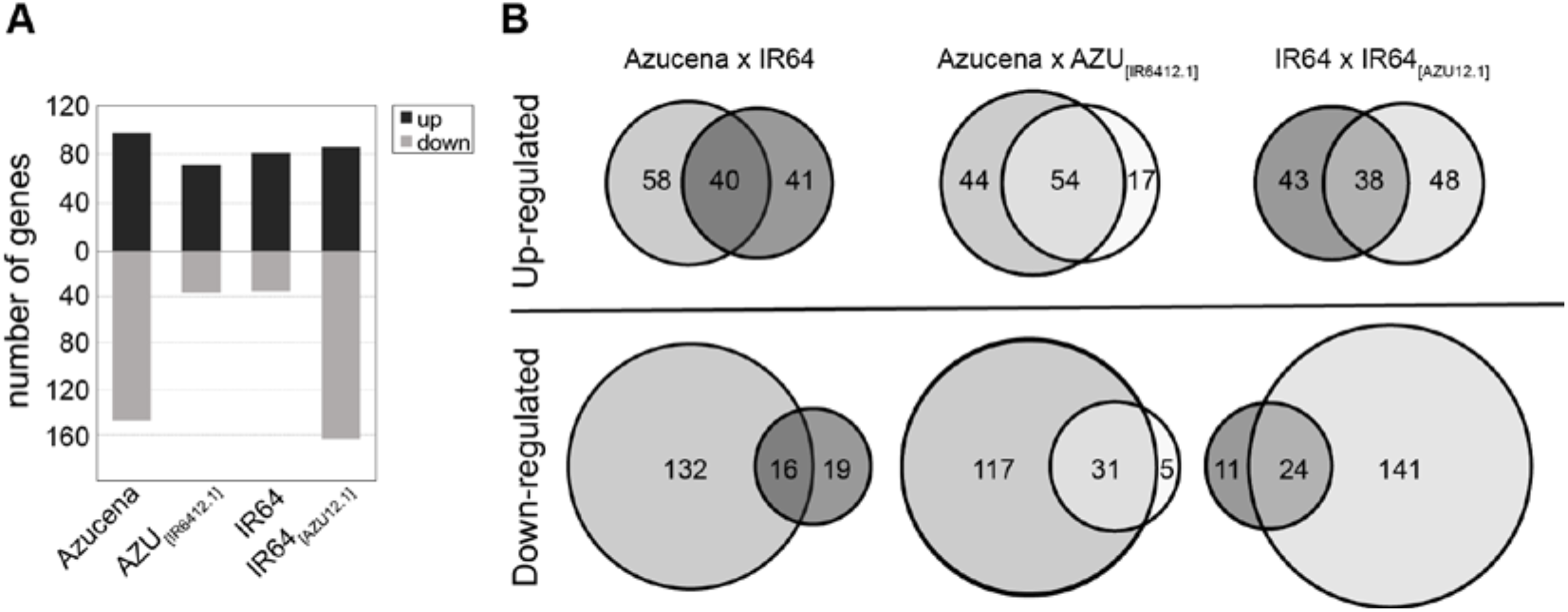
Azucena, IR64 and the reciprocal NILs AZU_[IR6412.1]_ and IR64_[AZU12.1]_ differ in their gene expression responses to Al stress. **(A)** Bar plot displaying the number of differentially-regulated genes in response to Al in roots of Azucena, IR64, and the reciprocal NILs AZU_[IR6412.1]_ and IR64_[AZU12.1]_ according to RNA-seq. Plants were treated with 0 or 80 μM Al^3+^ activity for 4 hours. **(B)** Venn diagrams of differentially-regulated genes in response to Al in each genotype.

### The effect of the reciprocal *Alt12.1* QTL introgressions on gene expression

We first examined the RNA-seq results for each NIL alongside its recurrent parent grown under control conditions (absence of Al). When the NIL AZU_[IR64121]_ was compared to its recurrent parent Azucena, 62 differentially-expressed genes were detected (Supplemental File S2). Of these, 40 are located within the introgression from IR64 in AZU_[IR6412.1]_, and are interpreted to represent genotypic differences between *indica* and *japonica*. Three of them were also found to be differentially-regulated by Al in Azucena: two were down- and one was up-regulated (Suppl. File S2). When the NIL IR64_[AZU12.1]_ was compared to IR64 under control conditions, 14 differentially-expressed genes were identified. Seven of them are located within the 2.36 Mb introgression from Azucena in IR64_[AZU12.1]_. Of these 14 genes, two were up-regulated by Al in IR64 (Suppl. File S2). None of the differentially expressed genes detected under control conditions corresponds to genes reported in previous studies to be Al-responsive (Yamaji et al., 2009; Tsutsui et al., 2012; Arenhart et al., 2014).

One gene, LOC_Os12g07340, located within the 44 Kb fine-mapped region of the *Alt12.1* QTL, was differentially expressed in both Azucena and IR64 backgrounds. According to RNA-seq, LOC_Os12g07340 expression is higher in Azucena than in the NIL AZU_[IR6412.1]_ (Supplemental Figure S4A) and also higher in the NIL IR64_[AZU12.1]_ than in IR64. This suggests that the Azucena allele is more highly expressed than the IR64 allele. Under Al stress, LOC_Os12g07340 is down-regulated by Al in lines carrying the Azucena allele, though the difference is only significant in Azucena (Suppl. Fig. S4B). LOC_Os12g07340 encodes an expressed protein with no known functional domains; whether this gene also contributes to the Al resistance phenotype associated with the *Alt12.1* QTL merits further investigation.

### The effect of the *Alt12.1* QTL introgressions on the transcriptional response to Al stress

While our RNA-seq data showed that the reciprocal *Alt12.1* introgressions triggered few changes in gene expression in the absence of Al, significant changes were observed in response to Al stress. In the Azucena genetic background, the presence of the *ART1* allele from IR64 reduced the number of up-regulated genes in the NIL AZU_[IR6412.1]_ (n = 71) relative to Azucena (n = 98). Of these, 54 were shared between Azucena and the NIL (Fig 4B). In contrast, in the IR64 genetic background the presence of the *ART1* allele from Azucena did not greatly affect the number of up-regulated genes (n = 81 in IR64 *versus* n = 86 in the NIL IR64_[AZU12.1]_), and 38 of these were shared between the two genotypes (Fig 4B).

The down-regulation of genes in response to Al stress showed a distinct pattern: 148 genes were down-regulated in Azucena, while only 36 were down-regulated in the NIL AZU_[IR6412.1]_ (Fig 4). Conversely, 35 genes were down-regulated in IR64, and 165 in the NIL IR64[azu_121_]. In other words, genotypes carrying the *ART1* allele from Al-resistant Azucena showed a greater number of down-regulated genes than those carrying the IR64 *ART1* allele. Of the 148 genes down-regulated in Azucena and 165 down-regulated in the NIL IR64_[AZU12.1]_, only 45 are in common. A GO analysis showed that the down-regulated genes in all four genotypes were strongly enriched in the categories ‘response to stimulus’ and ‘primary metabolic processes’ (Supplemental File S3). Azucena and both NILs were also enriched for genes involved in ‘signal transduction’ and ‘developmental processes’, but this was not true for IR64. The GO analysis suggests that the larger number of down-regulated genes in genotypes carrying the Azucena *ART1* allele encompasses a number of different functional categories (Supplemental Figure S5).

A comparison of global transcriptional responses between each NIL and its parent indicated that the reciprocal *Alt12.1* introgressions influenced the magnitude of the transcriptional response to Al. The average fold-change across all up-regulated genes was significantly lower in the NIL AZU_[IR6412.1]_ than in Azucena (Figure 5A, C). The same was true for down-regulated genes, which were significantly less down-regulated in the NIL AZU_[IR6412.1]_ than in the parent. Conversely, the NIL IR64_[AZU12.1]_ showed greater fold-changes in both up- and down-regulated genes than IR64 (Fig. 5B, D). These results suggest that genes in both genetic backgrounds are transcriptionally more responsive to the *ART1* allele from Al-resistant Azucena.

**Figure 5:**
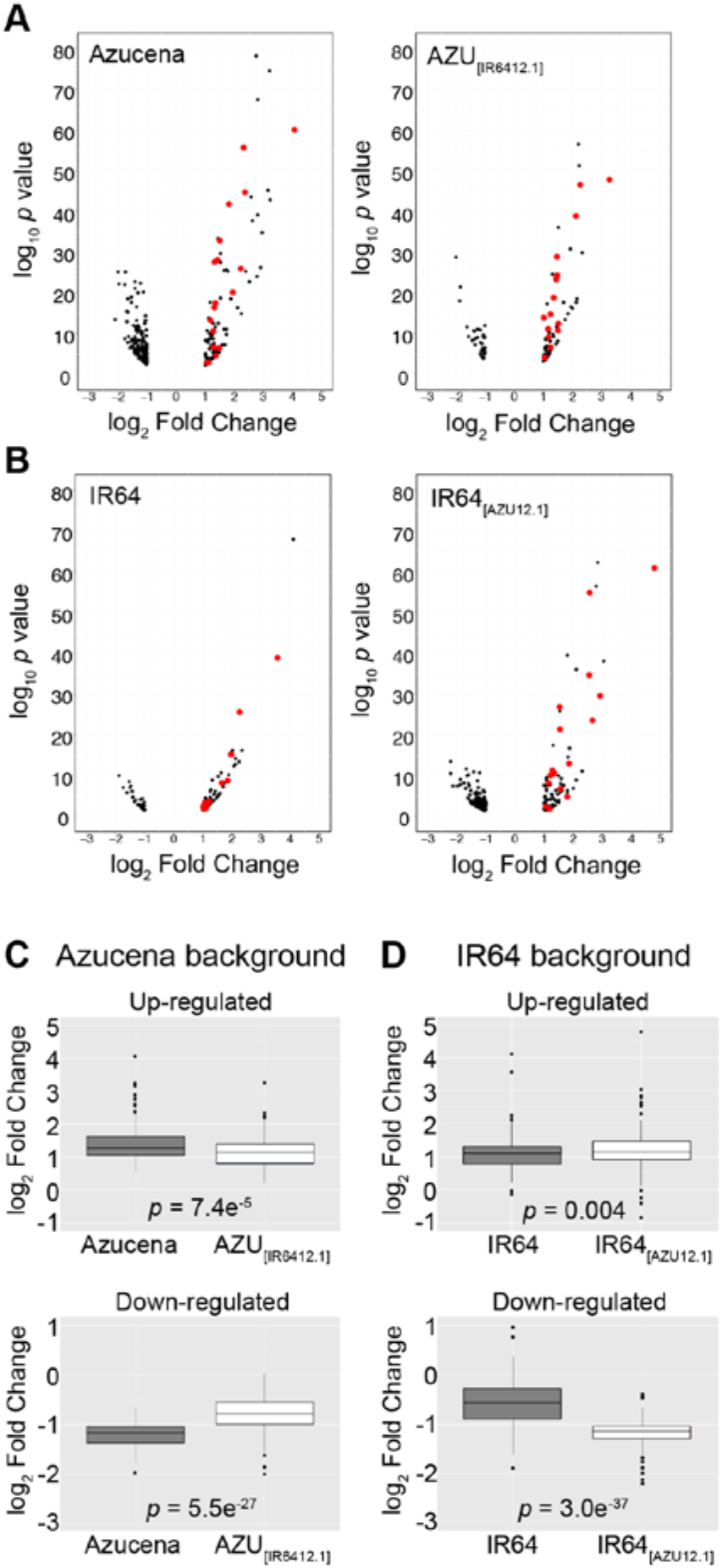
Gene expression responses to Al stress differ in magnitude in Azucena (*tropical japonica*) and IR64 (*indica*) genetic backgrounds. **(A)** and **(B)**: Volcano plots showing gene expression responses to Al stress in each genotype. The (-log_10_-transformed) *p* values were plotted against log_2_ fold-change for each differentially-regulated gene. **(A)** Azucena background; **(B)** IR64 background. Genes described by Yamaji et al. (2009) are highlighted in red. **(C)** and **(D)**: boxplots of gene expression changes (log_2_ fold change) of genes in Azucena **(C)** and IR64 **(D)** backgrounds. Genes up-regulated in Azucena in response to Al (N = 98) are significantly more up-regulated than those in the NIL AZU_[IR6412.1]_ (N = 71) (*p* value = 7.4e-05). Genes down-regulated in Azucena (N = 148) are significantly more down-regulated than those in the NIL AZU_[IR6412.1]_ (N = 36) (*p* value = 5.55e-27). Conversely, genes in IR64 (N = 81) are less up-regulated in response to Al than those in the NIL IR64_[AZU12.1]_ (N = 115) (*p* value = 0.004). Genes down-regulated in IR64 (N = 35) are significantly less down-regulated than those in the NIL IR64_[AZU12.1]_ (*p* value = 3e-37) (N = 164).

Next, we identified specific genes displaying higher fold-change in response to Al in the lines carrying the Azucena *ART1* allele (Figure 6). The heat map in Fig. 6A displays differentially-regulated genes in Azucena background, sorted according to the difference in fold-change in Azucena (native Azucena *ART1* allele) relative to the NIL carrying the IR64 *ART1* allele. Among those up-regulated by Al, 26 genes were > 1.5-fold more up-regulated in Azucena than in AZU_[IR6412.1]_. Of these genes, 12 were more than 2-fold more up-regulated in Azucena than in the NIL AZU_[IR6412.1]_ (Table I and Supplemental File S4).

**Figure 6:**
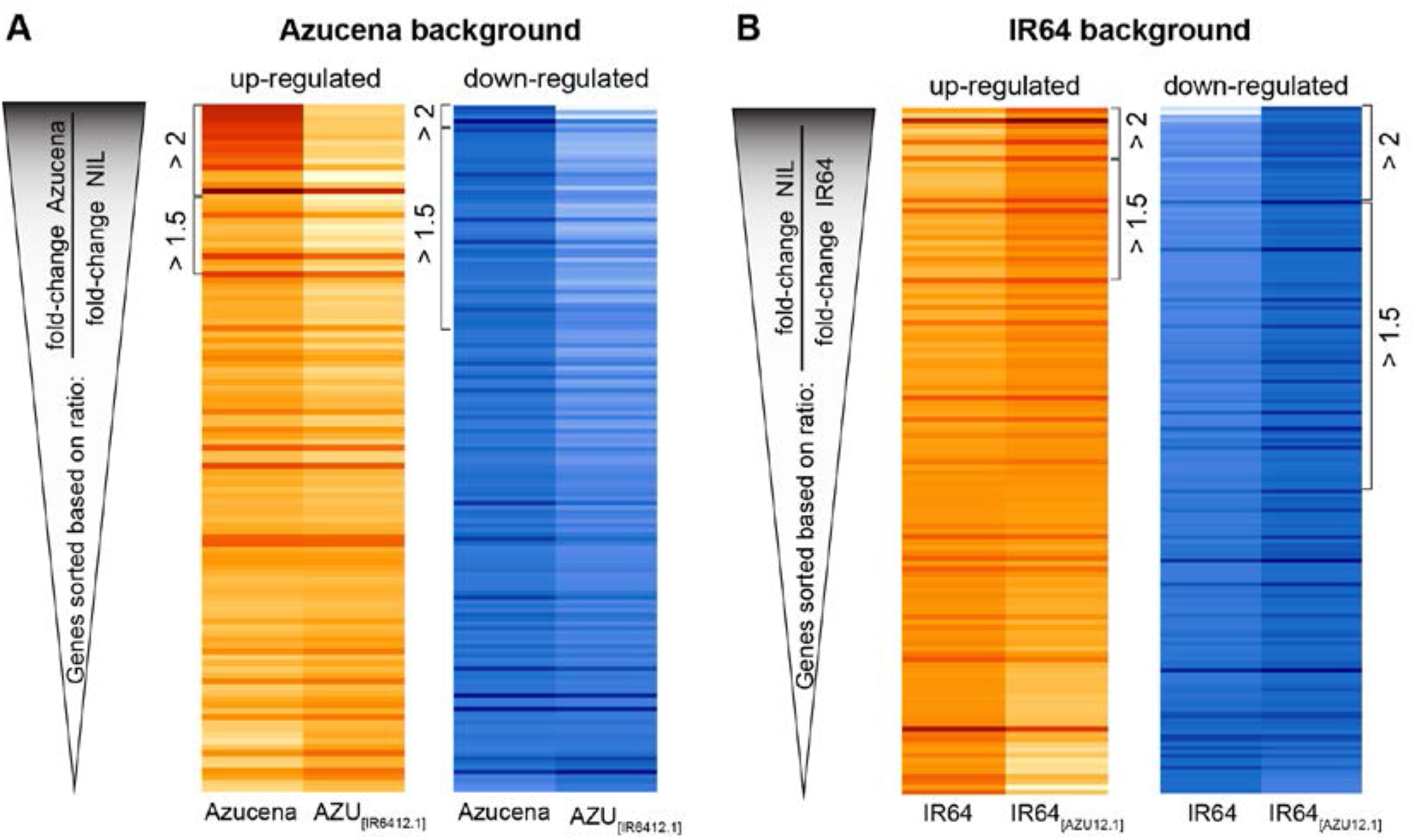
Differential gene expression responses to Al stress are stronger in the presence of the ART1 allele from Azucena in both genetic backgrounds. Heat maps illustrating the RNA-seq expression profile (log_2_ fold change) of differentially-regulated genes in response to Al in **(A)** Azucena and **(B)** IR64 backgrounds. Genes were sorted according to the ratio between the fold-change (in response to Al) in the line carrying the Azucena ART1 allele and the line carrying the IR64 ART1 allele. In other words, Azucena background lines were sorted based on (fold-change in Azucena / fold-change in NIL AZU_[IR6412.1]_). In the case of IR64 background, lines were sorted based on (fold-change in IR64_[AZU12.1]_ / fold-change in IR64). Brackets in **(A)** indicate genes with (fold-change Azucena / fold-change NIL AZU_[IR6412.1]_) ratios above 2 and above 1.5. Brackets in **(B)** indicate genes with (fold-change in IR64_[AZU12.1]_ / fold-change in IR64) above 2 and above 1.5. The color key represents normalized, log_2_-transformed fold-change. For up-regulated genes, red indicates high fold-change, light yellow indicates low (or no) fold-change. For down-regulated genes, dark blue indicates high fold-change; light blue indicates low (or no) fold-change.

**Table I:**
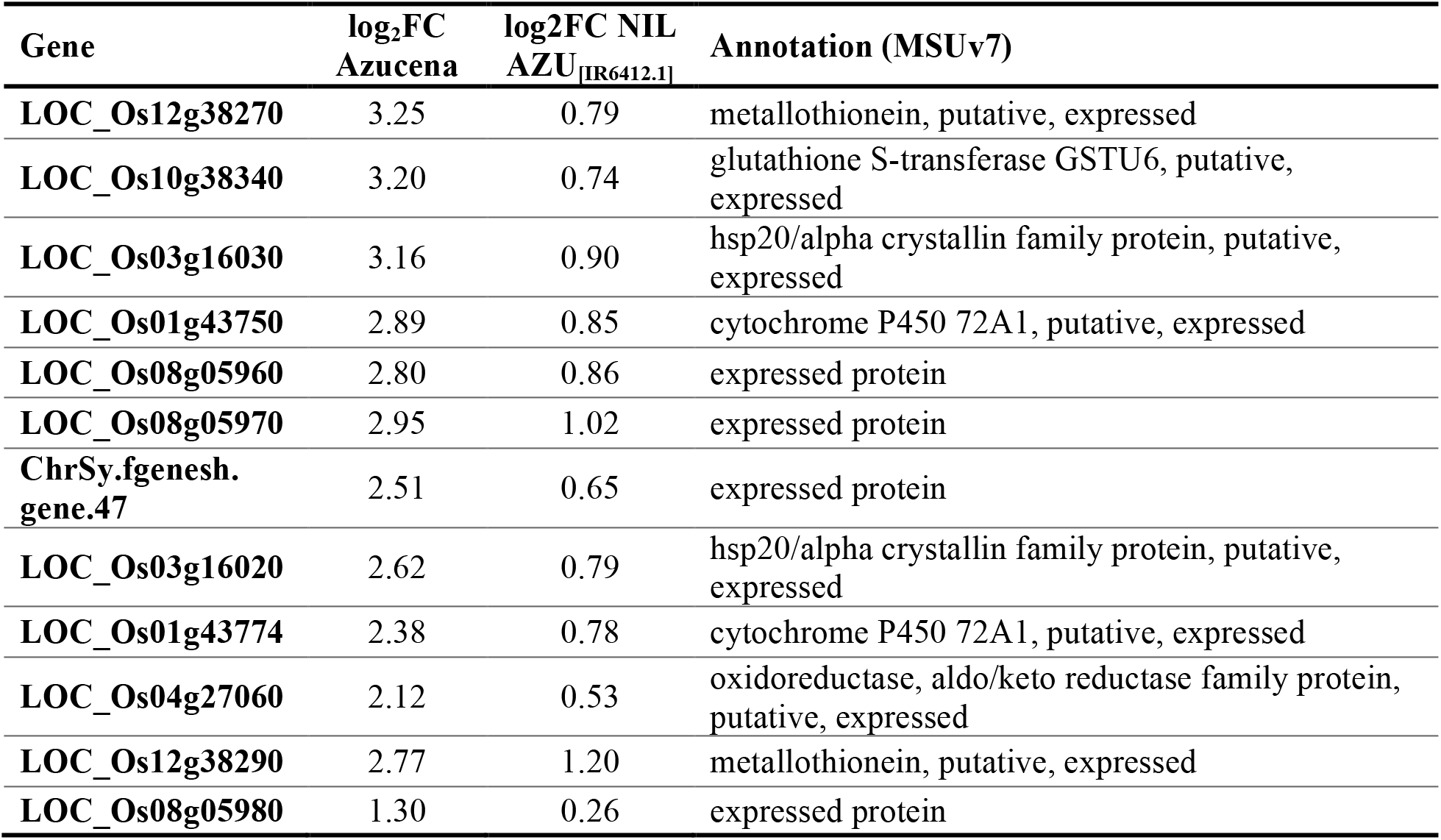
Genes up-regulated by Al stress showing fold-change 2X or higher in Azucena compared to NIL AZU_[IR6412.1]_.

These genes are at the top of the heat map (Fig. 6A). A similar pattern was observed for down-regulated genes. There were no genes displaying the opposite behavior, in other words, all differentially-regulated genes were either similarly or more intensely regulated in Azucena than in the NIL carrying the IR64 allele of *ART1*.

In the IR64 background (Fig. 6B), genes were sorted according to the difference in fold-change in the NIL IR64_[AZU12.1]_ (*ART1* allele from Azucena) relative to IR64, so that genes that respond more strongly in the presence of *ART1* from Azucena are at the top of the heat maps. Among the up-regulated genes, 32 genes were > 1.5-fold more up-regulated in IR64_[AZU12.1]_ than in IR64. Of these, ten were more than 2-fold more up-regulated in IR64_[AZU12.1]_ than in IR64 (Table II and Suppl. file S4). A number of genes were more up-regulated in IR64 than in the NIL IR64_[AZU12.1]_ (Suppl. File S4; bottom of heat map in Fig. 6B). The biological basis for this is not known; a possible explanation may be that IR64, being less Al-resistant, is under more stress and therefore more stress-related genes are differentially regulated.

**Table II:** Genes up-regulated by Al stress showing fold-change 2X or higher in the NIL IR64_[AZU12.1_] compared to IR64.

| Gene | log<sub>2</sub>FC<br>IR64 | log<sub>2</sub>FC NIL<br>IR64<sub>[AZU12.1]</sub> | Annotation (MSUv7) |
| --- | --- | --- | --- |
| LOC_Os06g19130 | 0.88 | 2.31 | cadmium tolerance factor, putative, expressed* |
| LOC_Os05g11320 | -0.13 | 1.16 | metallothionein-like protein 3B, putative, expressed |
| LOC_Os01g69010 | 3.56 | 4.79 | MATE efflux protein, putative, expressed (FRDL4)** |
| LOC_Os04g01690 | 0.65 | 1.87 | pyridoxal-dependent decarboxylase protein, putative, expressed |
| LOC_Os05g33900 | -0.04 | 1.09 | auxin-induced protein 5NG4, putative, expressed |
| LOC_Os12g38290 | 0.58 | 1.66 | metallothionein, putative, expressed |
| LOC_Os04g41750 | 1.85 | 2.92 | expressed protein** |
| LOC_Os11g41840 | 0.27 | 1.28 | transporter family protein, putative, expressed |
| LOC_Os06g39700 | 0.21 | 1.21 | DNA-directed RNA polymerase subunit alpha, putative, expressed |
| LOC_Os02g09390 | 1.65 | 2.65 | cytochrome P450, putative, expressed** |

\*gene annotation not confirmed by BlastP (see Supplemental File S4)
\*\*gene also listed by Yamaji et al. (2009)

Next, we examined whether the genes responding differently to the Azucena *versus* the IR64 *ART1* allele were the same or different in the two genetic backgrounds. A total of 26 genes showed greater up-regulation by the Azucena *ART1* allele in the Azucena background, while 32 showed greater up-regulation by the Azucena *ART1* allele in the IR64 background. Only six were shared across the two (Supplemental Figure S6A). A total of 48 genes showed greater down-regulation by the Azucena *ART1* allele in the Azucena background, while 95 showed greater down-regulation by the Azucena *ART1* allele in the IR64 background. 0nly eight were shared (Suppl. Fig. S6A). Although only two genotypes were tested, these results suggest that the genetic background influences the ART1-mediated transcriptional responses to Al stress. These results also indicate that many Al-responsive genes are independent of the *ART1* allele, or independent of ART1 altogether.

### Functional annotation of genes that respond more to Al stress in the presence of the Azucena *ART1* allele

In previous mutant studies, 32 genes were reported to be up-regulated by Al stress in wild-type but not in the *art1* mutant (Yamaji et al., 2009; Yokosho et al., 2011); these 32 genes have been designated ART1-regulated genes. Several of these were also identified in our study. The parental lines Azucena (Al-resistant) and IR64 (Al-sensitive) up-regulated 17 and 10 ‘ART1-regulated’ genes, respectively (Fig. 5). Of these, 8 were shared across the two genotypes (Suppl. Fig. S6B). The NIL AZU_[IR6412.1]_ up-regulated 14 genes; of these, 13 were shared between Azucena and the NIL. In the IR64 genetic background, the NIL IR64_[AZU12.1]_ up-regulated 17 ‘ART1-regulated’ genes while IR64 up-regulated 10, all of which were also up-regulated in the NIL.

The genes that respond more strongly to the *ART1* allele from Azucena are presumably the drivers of the phenotypic difference in Al resistance observed between NILs and their corresponding parents. Of the 26 genes more up-regulated in Azucena than in AZU_[IR6412.1]_, only three were previously identified as ART1-regulated genes (Suppl. File S4). Of the 32 genes that were more up-regulated in IR64_[AZU12.1]_ than in IR64, seven were previously listed as ART1-regulated (Suppl. File S4). These results suggest that in response to Al, ART1 regulates the expression of more genes than previously thought.

The previously described ART1-regulated gene *OsFRDL4* was more strongly up-regulated in the presence of the Azucena *ART1* allele than the IR64 allele in both Azucena and IR64 backgrounds (Suppl. File S4). *OsFRDL4* encodes a MATE family transporter that mediates Al-activated citrate exudation from rice roots (Yokosho et al., 2011). In our RNA-seq study, the expression of *OsFRDL4* is affected by the presence of different *ART1* alleles, with the Azucena *ART1* allele driving greater up-regulation under Al stress than the IR64 allele. Specifically, the expression of *OsFRDL4* is more up-regulated in response to Al in the NIL IR64_[AZU12.1]_ than in its recurrent parent IR64, and less up-regulated in response to Al in the NIL AZU_[IR6412.1]_ than in Azucena (Supplemental Figure S7, Suppl. File S4). The difference in *OsFRDL4* expression is associated with the presence of a 1.2-Kb insertion in the promoter of Azucena compared to IR64, suggesting that, in the genetic backgrounds used our study, variation in both the ART1 protein and the cis-regulatory region of *OsFRDL4* may contribute to the regulation of *OsFRDL4* expression in response to Al (Suppl. Fig. S7).

### ART1 *cis* element analysis

The *cis* element binding site of ART1 has been experimentally defined as the short, degenerate sequence motif GGN(T/g/a/C)V(C/A/g)S(C/G). This motif was identified based on protein gel-shift assays using the promoter of *STAR1* (Tsutsui et al., 2011). We set out to determine whether the putative regulatory regions of the Al-regulated genes in our whole-transcriptome analysis are enriched for the ART1-binding motif. As the first step, we probed the prevalence of the motif in the background, i.e. throughout the putative regulatory regions of all rice genes, regardless of their expression pattern. We generated a library of putative regulatory regions for all rice gene models based on the rice reference genome assembly (MSUv.7), defined as the 2-Kb region upstream of the start codon of each gene. When the ART1 cis-acting element GGNVS was mapped to the library of putative regulatory regions, the element was found to be present in 55,552 out of 55,554 putative regulatory sequences, with an average of 30.3 motifs per 2 Kb sequence (Supplemental Figure S8A). Because ART1 was shown to bind with lesser affinity when certain bases were found in positions 3 and 4 of the motif (Tsutsui et al., 2011), we also mapped the less degenerate motif GGYMS, in which the bases predicted to have less affinity to ART1 were excluded. The motif GGYMS was found in 55,460 out of 55,554 putative regulatory sequences, with an average of 10 hits per sequence (Suppl. Fig. S8B). We determined that enrichment of the 5-bp ART1-binding *cis* element in the regulatory region of putative Al response genes is insufficient criteria for identifying ART1-regulated genes, due to the widespread presence of the motif in the background of putative regulatory regions.

### Putative ART1-regulated genes provide insight into transgressive segregation for Al resistance in rice

We identified a number of genes that responded to Al stress only in the IR64 (Al-sensitive parent) genetic background, including two previously identified as ART1-regulated genes by Yamaji et al. (2009): LOC_Os01g53090 and LOC_Os04g41750. In our RNA-seq dataset, these genes were significantly up-regulated in response to Al stress in both IR64 and the NIL IR64_[AZU12.1]_ but not in Azucena or AZU_[IR6412.1]_ (Figure 7A). In other words, these two genes were up-regulated only in the IR64 genetic background and not in the Azucena genetic background, and the up-regulation is triggered by either of the *ART1* alleles. We hypothesized that IR64, the Al-sensitive parent, carries positive alleles conferring enhanced Al resistance at both of these loci (and possibly others). In line with this hypothesis, we previously reported transgressive segregation for rice Al resistance (Famoso et al., 2011).

**Figure 7:**
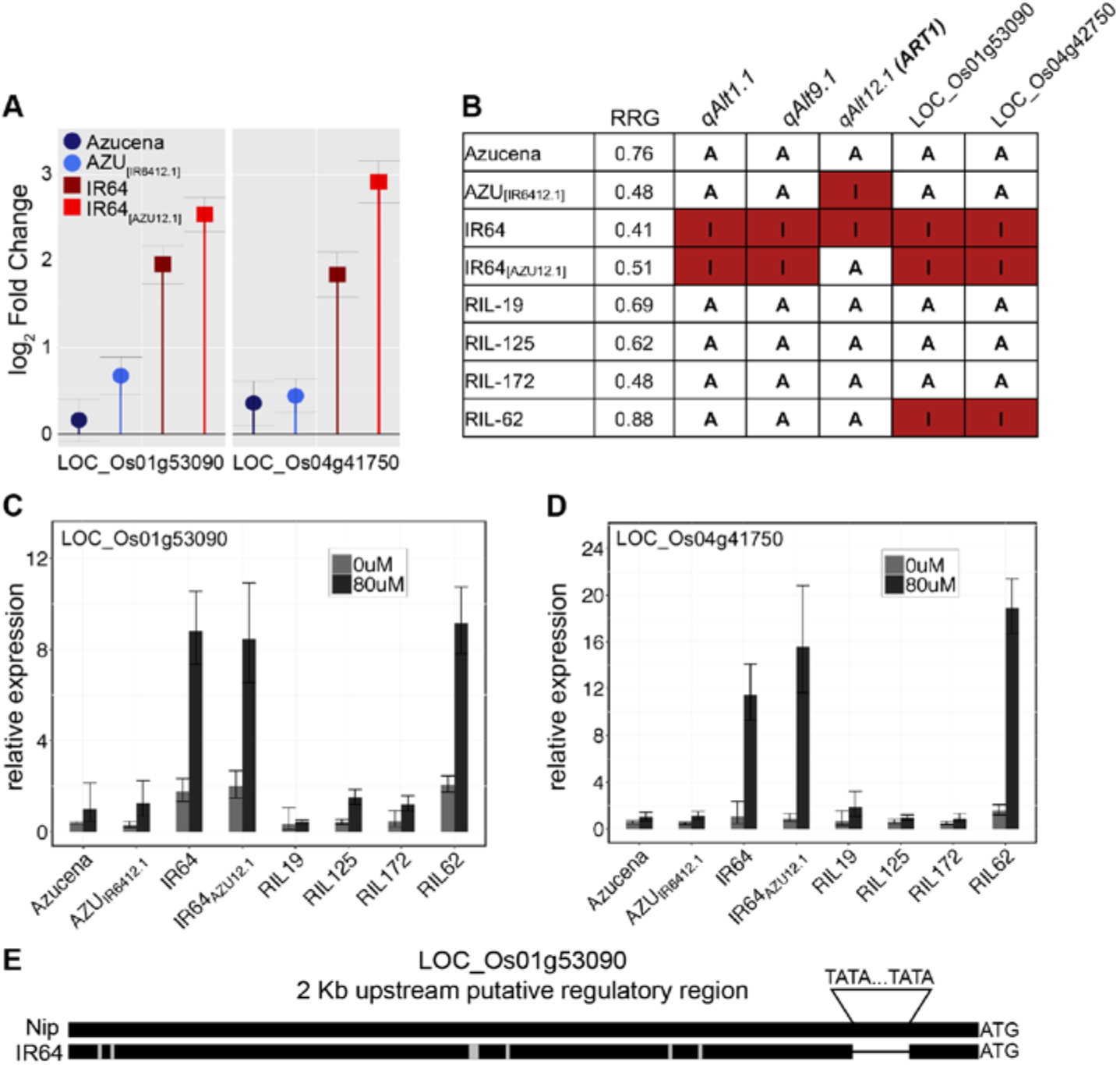
Expression patterns of ART1-regulated genes in different genetic backgrounds provide clues about transgressive variation. **(A)** Relative expression based on RNA-seq (log_2_ fold change) of two putative ART1-regulated genes, LOC_Os01g53090 and LOC_Os04g41750 under Al stress. **(B)** Genotypic makeup and Al resistance phenotype (RRG) of rice genotypes used in qRT-PCR study: the parents Azucena and IR64, the reciprocal NILs IR64_[AZU12.1]_ and AZU_[IR6412.1]_, and RILs 19, 125, 172 (controls) and 62. RIL-62 shows transgressive variation for Al resistance. The colored boxes indicate genotype at these loci: QTLs *Alt1.1, Alt9.1* and *Alt12.1*, as well as at the putative ART1-regulated genes LOC_Os01g53090 and LOC_Os04g41750. “A” indicates Azucena genotype at that given loci; red boxes with “I” indicate IR64. **(C)** and **(D)** Relative expression based on qRT-PCR for genes **(C)** LOC_Os01g53090 and **(D)** LOC_Os04g41750 in roots of Azucena, AZU_[IR6412.1]_, IR64, IR64_[AZU12.1]_, RIL-19, RIL-125, RIL-172, and RIL-62 grown in hydroponics and treated for 4h under control (0 μM Al^3+^) (light grey bar) and Al stress conditions (80 μM Al^3+^ activity) (dark gray bar). Error bars indicate a 95% confidence interval. **(E)** Graphical representation of a DNA sequence alignment of the 2Kb putative regulatory region of LOC_Os01g53090 between the reference genome (Nipponbare) and IR64. Black bars represent regions of sequence identity. Grey rectangles indicate small indels and SNPs. A bracket indicates the position of a “TA” sequence repeat (32 bp longer in Nipponbare/Azucena relative to IR64).

To further explore this hypothesis, we investigated the expression patterns of LOC_Os01g53090 and LOC_Os04g41750 in a line (RIL-62) from the RIL population that carries IR64 alleles at both of these ART1-regulated loci; this RIL also carries favorable alleles from Azucena at all three previously reported Al resistance QTL (*Alt1.1, Alt9.1* and *Alt12.1*, i.e. *ART1*) (Famoso et al., 2011), and shows transgressive variation for Al resistance (Fig. 7B). As controls, we selected three additional RILs carrying Azucena (resistant) alleles at the three QTL, as well as at the two ART1-regulated loci: RIL-19, RIL-125 and RIL-172 (Fig. 7B). Using qRT-PCR, we confirmed the observations from RNA-seq showing that both genes were strongly up-regulated in response to Al in IR64 and IR64_[AZU12.1]_ but not in Azucena or AZU_[IR6412.1]_. Both genes were also strongly up-regulated by Al in the transgressive RIL-62, but not in any of the control RILs (Fig. 7C,D).

It is possible that variation in the *cis* regulatory regions of these genes may contribute to the difference in gene expression patterns. In fact, in the upstream regulatory region of LOC_Os01g53090 there is a “TA” repeat region just 97 bp upstream of the start codon. An alignment between the Nipponbare reference genome and the IR64 assembly showed that the repetitive region is 32 bp longer in Nipponbare (Fig. 7E). LOC_Os04g41750 is not only up-regulated exclusively in the IR64 background, but also responds more strongly to Al in the presence of the ART1 from Azucena: the up-regulation of LOC_Os04g41750 in response to Al in the NIL IR64_[AZU12.1]_ is higher than in IR64 (Table II and Fig. 7). These results may illustrate how recombination between divergent parents can bring together favorable genetic variation at interacting loci that contribute to transgressive phenotypic variation in the offspring.

## Discussion

### Allelic variation in *ART1* affects rice Al resistance

Our present study focused on a major QTL for Al resistance in rice, *Alt12.1*. This QTL was identified in an RIL mapping population derived from a cross between an Al-resistant *tropical japonica* variety (Azucena) and an Al-sensitive *indica* variety (IR64). Via fine-mapping, we narrowed down the QTL *Alt12.1* to a region of 44 Kb surrounding *ART1* (Fig. 2). *ART1* encodes a C2H2 transcription factor required for rice Al resistance, and regulates the expression of other known Al resistance genes. While we did not observe differences in *ART1* transcript levels between Azucena, IR64, or the NILs (with or without Al treatment), we identified extensive sequence polymorphism in the coding region. These polymorphisms are not predicted to truncate the protein product or affect the C2H2 domain but could affect protein folding and interaction with target gene promoters. It is interesting to note that most of the polymorphisms between *indica* and *japonica ART1* alleles occur at the C-terminal end of the protein. Perhaps the most notable polymorphism is the large C-terminal insertion present in Kasalath and IR64, and absent in both Azucena and Nipponbare (Fig. 2). This insertion adds a poly-threonine sequence downstream of a polyproline-like sequence. The reactive hydroxyl groups on threonine are common targets for posttranslational modifications making this an interesting polymorphism for future protein-protein interaction studies.

The RNA-seq results presented here indicate that the *ART1* allele from Azucena leads to a stronger transcriptional response to Al stress in rice roots. Using yeast one-hybrid assays, we demonstrate that the ART1 proteins encoded by the alleles found in *indica* / *aus* (IR64; Kasalath) and *japonica* (Azucena; Nipponbare) differ significantly in their DNA binding ability. In the yeast one-hybrid, the ART1 proteins encoded by the IR64 and Kasalath alleles activated the reporter gene more strongly than the ones from Azucena and Nipponbare (Fig. 3C). This is a non-intuitive result, given that the Azucena allele drives gene expression more strongly than the IR64 allele *in planta*. Yeast one-hybrid is a heterologous system, in which the TF of interest is fused with the strong GAL4 activation domain and the DNA sequence of interest is placed upstream of a reporter gene. As such, other elements that may affect TF activity *in planta* are missing. Further studies using promoter-reporter systems in plant protoplasts will shed more light into the transcriptional activation ability of the different ART1 proteins.

### Using reciprocal NILs to study transcriptomic responses to Al in IR64 and Azucena

We generated reciprocal NILs in which the *Alt12.1* QTL region from Azucena was introgressed into IR64, and vice-versa (Fig 1). The NILs validated the effect of the QTL on Al resistance in both genetic backgrounds: the NIL in the Azucena background, AZU_[IR6412.1]_, is less Al-resistant than Azucena; conversely, the NIL in the IR64 background, IR64_[AZU12.1]_, is more Al-resistant than IR64. Because ART1 is a transcription factor, the phenotypic differences observed between each NIL and their respective recurrent parent are likely to result from differences in the expression of genes regulated by ART1. We took advantage of the reciprocal NILs to study how the IR64 and Azucena *ART1* alleles affect transcriptomic responses to Al stress in both genetic backgrounds.

We performed RNA-seq in roots of rice seedlings of Azucena, IR64 and the reciprocal NILs AZU_[IR6412.1]_ and IR64_[AZU12.1]_ under control conditions (no Al) or exposed to Al for 4 hours. To determine the effect of the *Alt12.1* QTL introgressions on gene expression under control conditions, we compared gene expression in each NIL and its recurrent parent in the absence of Al. Under these conditions, only a small number of differentially-regulated genes were identified. The majority localized to the chromosomal introgressions in each of the NILs, suggesting that these genes represent “background” genotypic differences between *indica* and *japonica*. However, five genes showed differential expression in response to Al treatment (Suppl. File S2). This included LOC_Os12g07340, which is one of six genes located within the fine-mapped region, only 28 Kb away from *ART1* (Fig. 2A). The function of these genes is unknown, and though none are predicted to encode a regulatory protein or TF, we cannot rule out the possibility that one or more may contribute to Al resistance. Our study illustrates some of the complexities and limitations of using NILs to molecularly characterize genes underlying QTLs. Even if the background of the NIL is clean, the target introgressions inevitably harbor more than just the gene of interest. Genome-editing tools are now well-established in a variety of plant species; we are currently applying this technology to create “clean” allelic swaps for *ART1* in both *indica* and *japonica* backgrounds.

### Down-regulation of gene expression in response to Al is associated with the Azucena *ART1* allele

One of the most striking features of our RNA-seq study is the extent of the down-regulation response to Al stress in Azucena when compared to IR64 (Fig. 4). Moreover, the larger number of down-regulated genes is associated with the presence of the Azucena *ART1* allele: Azucena and the IR64 NIL carrying the *Alt12.1* introgression from Azucena (IR64_[AZU12.1]_) both displayed a larger number of down-regulated genes than the genotypes carrying the *ART1* allele from IR64 (IR64 and AZU_[IR6412.1]_). Based on these results as well as the Y1H findings, it is tempting to speculate that ART1 may act as a negative regulator of Al responses. However, the role of ART1 as an activator of transcription is well established, as the literature provides many examples of genes involved in Al resistance in rice that are up-regulated by the stress and require ART1 for their activation. However, the possibility exists that ART1 may interact with cis-regulatory elements and other transcription factors to act as both an activator and a repressor of transcription.

To date, most transcriptomics studies examining responses to Al stress have focused on up-regulated genes. For instance, Yamaji et al. (2009) did not report on down-regulated genes in the *art1* mutant study, so it is not known whether the *art1* mutant also displayed less gene down-regulation in response to Al. Transcriptome studies in the Al-sensitive species *Medicago truncatula* (Chandran et al., 2008) and in the highly Al-resistant species buckwheat (Yokosho et al., 2014) did not observe a larger number of down-regulated versus up-regulated genes. In maize, transcriptional profiling comparing an Al-resistant and an Al-sensitive genotype (Maron et al., 2008) reported that the Al-sensitive genotype showed a larger number of both up- and down-regulated genes. This is likely due to higher stress levels resulting from Al toxicity, as the number of differentially-regulated genes in the Al-sensitive genotype increased over time. In our study in rice, the Al-resistant genotype displayed a larger number of down-regulated genes. However, we can only draw limited conclusions from these studies, as each study analyzed a small number of lines. More research is needed before we can implicate gene down-regulation with specific mechanisms of Al resistance.

### The effect of *ART1* on transcriptomic responses to Al stress may depend on the genetic background

Our RNA-seq study identified a number of differentially-regulated genes in response to Al stress, and only about half of them were shared between the Azucena (*tropical japonica*) and IR64 (*indica*) genetic backgrounds. The ART1 from Al-resistant Azucena led to greater global changes in gene expression in response to Al stress when compared to the ART1 from IR64, both in the number of genes and magnitude of the response. This result was observed in both the Azucena (native) and the IR64 genetic background (NIL IR64_[AZU12.1]_, carrying the *Alt12.1* QTL from Azucena), suggesting that the Azucena *ART1* allele is more effective in conferring Al resistance than the IR64 allele in both genetic backgrounds. The small sample size (n = 4 genotypes) limits our ability to statistically analyze the differences between genetic backgrounds; nevertheless, our data suggests that the identity of the genes affected by the two *ART1* alleles differed in Azucena and IR64.

### ART1 allele swapping between Azucena and IR64 affects the expression pattern of many genes not previously designated as ART1-regulated

Among the genes up-regulated by Al stress in our RNA-seq study are many previously characterized Al resistance genes, including *OsNrat1, OsFRDL4, OsFRDL2, OsALS1, STAR1, STAR2* and *OsCDT3. OsNrat1* encodes a plasma membrane-localized Al uptake transporter (Xia et al., 2010; Li et al., 2014). *OsNrat1* function is likely coupled with a mechanism of internal detoxification, involving another ART1-regulated gene: *OsALS1*. This gene encodes a half-size ABC transporter localized to the tonoplast of root cells, where it is thought to sequester Al^3+^ into the vacuole (Huang et al., 2012). *STAR1* and *STAR2* were also shown to play a role in rice Al resistance: *STAR1* encodes the nucleotide-binding domain of a bacterial-type ABC transporter that interacts with the transmembrane domain encoded by *STAR2* (Huang et al., 2009). Disruption of either gene results in higher Al sensitivity; however, the mechanism by which the STAR1-STAR2 complex confers Al resistance is unknown. When expressed in *Xenopus* oocytes, STAR1-STAR2 facilitates the export of UDP-glucose; this substrate is proposed to modify the cell wall in a way that reduces Al^3+^ accumulation. *OsCDT3* encodes a small cysteine-rich peptide that exhibits Al-binding properties (Xia et al., 2013).

The genes that respond more strongly to the *ART1* allele from Azucena than to the *ART1* allele from IR64 are presumably the drivers of the phenotypic difference in Al resistance observed between NILs and their corresponding recurrent parents. Among the genes previously designated as ART1-regulated, only a few were among those that responded more strongly to the *ART1* allele from Azucena: *OsFRDL4*, LOC_Os02g51930 and LOC_Os02g09390 in the Azucena background, and *OsFRDL4*, LOC_Os04g41750, LOC_Os02g09390, LOC_Os11g29780, LOC_Os10g38080, LOC_Os03g55290 and LOC_Os09g30250 in the IR64 background (Suppl. File S4). The functions of the vast majority of genes that responded more strongly to Al stress in the presence of the *ART1* allele from Al-resistant Azucena are unknown and present new opportunities for exploring the mechanisms of transcriptional regulation in rice roots in response to Al stress.

### Expression of the MATE transporter *OsFRDL4* is dependent on genetic background

*OsFRDL4* encodes a Multidrug and Toxin Extrusion (MATE) transporter that mediates root citrate release (Yokosho et al., 2011); Al-activated release of organic acids from the root is a major physiological mechanism of plant Al resistance (Kochian et al., 2015). *OsFRDL4* is the major candidate for the gene underlying an Al resistance QTL in a Nipponbare (*temperate japonica*) x Kasalath (*aus*) population (Yokosho et al., 2016). The authors of that study also reported a 1.2-Kb insertion in the promoter of *OsFRDL4*, and demonstrated that a majority of *japonica* varieties tested carried the insertion, while most *indica* varieties did not, and suggested that the insertion was associated with higher levels of *OsFRDL4* expression under Al stress. Yokosho et al. (2006) also concluded that the promoter of *OsFRDL4* responds equally to the ART1 proteins from Nipponbare and Kasalath. In contrast, our RNA-seq study indicates that the expression of both alleles of *OsFRDL4* is affected by the presence of the different *ART1* alleles from Azucena and IR64, with the Azucena ART1 driving greater up-regulation than the IR64 ART1. In the Azucena background, *OsFRDL4* carries the 1.2-Kb insertion in the promoter (like Nipponbare), while IR64 does not (like Kasalath), and in both cases, *OsFRDL4* expression was more up-regulated by the Azucena ART1 allele than by the IR64 ART1 allele. It is possible that other regulatory elements are involved in regulating *OsFRDL4* expression in response to Al. Similarly, (Melo et al., 2013) reported that sorghum NILs carrying different alleles of the major Al resistance gene *SbMATE* exhibited only partial transfer of Al resistance, which was closely correlated with a reduction in *SbMATE* expression. The authors suggest that *SbMATE* expression is regulated by multiple elements, with the relative importance of each element depending on genetic context. Our results suggest that this may also be the case for *OsFRDL4* expression in rice. Moreover, these results further illustrate the importance of genetic background, not only when selecting alleles for breeding purposes, but also when assessing gene function at the molecular level.

### ART1 *cis* element

The binding site of ART1 was experimentally defined as the short, degenerate sequence motif GGNVS, found in the promoter region of genes described as ART1-regulated. This motif was further validated for *OsSTAR1*, a known target of ART1, using EMSA (Tsutsui et al., 2011; Yokosho et al., 2011). Our results show that the presence of the *cis* element alone does not provide evidence of ART1 binding, as the short binding site is present in the promoter of almost all predicted rice genes. The very high frequency of the ART1 *cis* element in upstream regulatory regions of rice genes is not unexpected. Because the motif is both very short and very degenerate, it has a high probability of being found by chance. Evidence from the literature suggests that plant *cis* elements defined based on *in vitro* DNA-binding studies often contain core binding sites that are short, degenerate, and shared by multiple TFs within a transcription factor gene family (see AtcisDB: http://arabidopsis.med.ohio-state.edu/AtcisDB, and Rombauts et al., 2003). Several mechanisms control TF binding to a *cis* element only in a specific context, including sequence specificity of the TF, combinatorial cooperation between TFs, and chromatin state (Vandepoele et al., 2009; Vaahtera and Brosché, 2011; Franco-Zorrilla et al., 2014).

It is worth noting that the list of putative ART1 targets was originally based on genes significantly mis-regulated in the *art1* mutant background. While this is a common method for defining genes regulated by a specific transcriptional activator, it can overestimate the number of true targets as TFs often regulate other TFs in complex regulatory networks. Because many TFs have low-abundance transcripts, they frequently fall below standard cutoff values in microarray and RNA-seq analyses. Thus, some of these genes may be indirectly regulated by ART1 via unidentified transcriptional regulators. Direct binding of the ART1 protein to upstream regulatory regions has only been experimentally demonstrated for *STAR1* (Tsutsui et al., 2011) and *OsFRDL4* (Yokosho et al., 2016). Furthermore, while the *art1* mutant phenotype is clearly severe, it is unclear whether it is a complete loss-of-function allele. While our own yeast one-hybrid work shows that the aberrant *art1* protein encoded by the *art1* mutant does not interact with the *OsSTAR1* promoter, our YFP fusion experiment suggests that the *art1* mutant protein does localize properly to the nucleus and contains at least one DNA-binding domain. Thus, we cannot rule out the possibility that the *art1* mutant protein disrupts the transcription of non-native ART1 targets. Nevertheless, data from our RNA-seq study of natural, functional variants suggests that ART1 has a distinct effect on transcription that is highly dependent on genetic background, and not necessarily all of the mis-regulated genes are directly regulated by ART1. Future work to validate bona-fide ART1 transcriptional targets, including TF network analysis using more in-depth transcriptome data, will likely improve our functional understanding of ART1 and help to better refine its cis-regulatory element.

### *ART1*-regulated genes and transgressive variation for rice Al resistance

The Azucena (*tropical japonica*) X IR64 (*indica*) mapping population exhibited transgressive segregation for Al resistance (Famoso et al., 2011). This phenomenon can occur when the susceptible parent (in this case the Al-sensitive *indica* IR64) contributes positive alleles to the transgressive offspring. Using RNA-seq, we identified a number of genes that responded to Al stress only in the IR64 genetic background, including two genes previously identified as ART1-regulated by Yamaji et al. (2009): LOC_Os01g53090 and LOC_Os04g41750. The functions of these genes are not known. A BlastP search revealed LOC_Os04g41750 encodes a protein containing two DUF642 domains; DUF642 proteins constitute a highly conserved family of proteins that are associated with the cell wall and are specific to spermatophytes (Vázquez-Lobo et al., 2012). Two Arabidopsis DUF642 proteins were recently shown to regulate the activity of pectin methyl-esterase during seed germination (Zúñiga-Sánchez et al., 2014). The cell wall plays an important role in rice Al resistance, but the exact mechanism(s) are far from understood. Therefore, the molecular function of LOC_Os04g41750 is worthy of further investigation.

### ART1 function beyond Al resistance?

While a majority of genes differentially expressed between each NIL and its parent under control conditions was localized to the chromosomal introgression, a small number of them were distributed on other chromosomes. Therefore, the possibility that ART1 also regulates gene expression in the absence of Al cannot be discarded. It is important to note that expression of *ART1* itself is constitutive and is not up-regulated by Al (Fig. 2C); the signaling mechanism that activates ART1 binding to the promoter of Al-responsive genes in response to Al stress has not been elucidated. In addition, it is also possible that ART1 could have additional functions beyond Al resistance. In fact, *SENSITIVE TO PROTONRHIZOTOXICITY1* (*STOP1*), the homolog of *ART1* in *Arabidopsis thaliana*, was recently implicated as a critical checkpoint in the root developmental response to phosphate starvation (Mora-Macías et al., 2017). STOP1 also regulates low pH and Al stress responses in Arabidopsis (Iuchi et al., 2007); therefore, the authors suggest that STOP1 is likely to orchestrate root responses to multiple environmental stresses. The *art1* mutant did not show decreased root elongation when exposed to cadmium, lanthanum, zinc or copper (Yamaji et al., 2009); however, it was not phenotyped under stresses that commonly occur together with Al as part of the acid soil syndrome, such as phosphate deficiency and iron toxicity. Plants are more likely to have evolved regulatory mechanisms that respond to multiple environmental stresses that are frequently found together in nature (Maron et al., 2016). Phenotyping of the reciprocal NILs carrying the *ART1* alleles from *indica* and *japonica*, as well as the *art1* mutant, under other abiotic stresses that co-occur in acid soils will shed light into the possible functions of ART1 beyond Al resistance. It will also deepen our understanding of how a TF such as ART1 can mediate quantitative forms of stress resistance, via transcriptional regulation of an orchestra of downstream genes, each with its own potential for variation that contributes to the fine-tuning of plant response to stress.

## Materials and Methods

### Plant Materials and Plant Growth Conditions

Seeds of the Al-resistant rice variety Azucena (*Oryza sativa* ssp. *tropical japonica*), of the Al-sensitive variety IR64 (*Oryza sativa* ssp. *indica*), the recombinant inbred lines RIL-241, and RIL-48 were originally obtained from the Institut de recherche pour le développement (IRD, Montpellier, France) but have been amplified in the Guterman Greenhouse at Cornell University. Experiments in hydroponic nutrient solutions under control (0 μM Al^3+^ activity), and Al stress (80 or 160 μM of Al^3+^ activity) were conducted according to Famoso et al. (2010).

### Phenotyping for Al resistance

Al resistance was phenotyped as described by Famoso et al. (2010). Individual root seedling digital images were obtained to quantify total root growth (TRG) values according to (Clark et al., 2013) using the software RootReader2D (www.plantmineralnutrition.net/rr2d.php). Total root growth (TRG) values from each genotype grown under control and Al stress conditions were used to estimate relative root growth (RRG) indices as described (Famoso et al., 2010). RRG values were used for further statistical analysis.

### Genotyping

For PCR analysis total DNA was extracted from fresh leaf tissue using the Extract-N-Amp^™^ Plant Kit according to manufacturer’s instructions (Sigma-Aldrich, www.sigmaaldrich.com/). Plants were genotyped using InDel and SNP markers based on competitive allele-specific PCR KASP^™^ chemistry assays (LGC, www.lgcgroup.com/). InDel and KASP^™^ SNP primers were identified and designed according to Imai et al. (2013) (Supplemental Table S2). InDel PCR was performed using 20 ng of DNA as template and amplified using GoTaq^®^ Green Master Mix (Promega, www.promega.com/). Conditions for amplification and visualization of InDel markers are described in Arbelaez et al. (2015). For the KASP^™^ markers 10 ng of DNA was used as template and the KASP^™^ Assay/Master mixes (LGC, www.lgcgroup.com/) were used to amplify the KASP^™^ PCR products. PCR conditions and graphical viewing of genotyped KASP^™^ markers were carried out as described by Imai et al. (2013). Samples were genotyped using the 6K Infinium array (Illumina, www.illumina.com/). DNA was extracted using a modified CTAB protocol described by (Imai et al., 2013). The 6K Infinium assays were performed as described by (Arbelaez et al. (2015).

### Quantitative real-time PCR (RT-qPCR)

Total RNA was isolated using the QIAGEN RNeasy Mini Kit according to manufacturer’s instructions (https://www.qiagen.com). In-column digestion of genomic DNA was performed using the QIAGEN RNase-Free DNase set according to manufacturer’s instructions. Singlestranded cDNA was synthesized from total RNA using the High Capacity RNA-to-cDNA Kit (Applied Biosystems) according to manufacturer’s instructions. Quantitative real-time PCR was performed employing the relative standard curve method using Power SybrGreen Mastermix (Applied Biosystems). Gene expression was normalized against three endogenous controls. Primers used were as follows. *ART1:* 5’-CCAGCCGCTGAAGACGAT-3’ and 5’-GCAGTGGCTCCGCTTGTAGT-3’; LOC_Os01g53090: 5’-CGTGAACAGCCTCTTCGAG-3’ and 5’-CCATCTCCCATGTCTTGATCG-3’; LOC_Os04g41750: 5’-ACTCGTTCGCCATCAAGAC-3’ and 5’-AGTAGTCGTCCTCGTCCATC-3’. Endogenous control genes: HNR, 5’-TAAGGTCGGTATCGCCAATC-3’ and 5’-GGCAGGTTCTGCAGTGGTAT-3’; Tip41, 5’-CGCTCCAGCTCTTTGAAGATAAA-3’ and 5’-ACTCTCCCCAAAAACCATCTCA-3’; 18S, 5’-GACTACGTCCCTGCCCTTTG-3’ and 5’-TCACCGGACCATTCAATCG-3’. Three technical replicates were averaged per sample per each assay. RNA from six independent biological replicates was used to measure *ART1* expression and confirm that there are no significant differences in expression between genotypes and/or treatments. One biological replicate was used to confirm LOC_Os01g53090 and LOC_Os04g41750 in NILs and parents, and to determine expression levels in the RILs.

### Subcellular localization of ART1:YFP protein fusions

In order to determine the cellular localization of the aberrant protein generated by the *art1* mutant, we re-created the mutation *in vitro.* This clone was also used to generate the yeast one-hybrid construct. To generate the mutant protein, we used the reference genome sequence to identify the new stop codon in the frameshifted protein, which extended into the 3’UTR of the *ART1* locus. We then used a series of long oligos to add the required nucleotides to the previously-cloned Nipponbare allele, followed by USER-based cloning to introduce the 1-bp deletion (http://www.cbs.dtu.dk/services/PHUSER/).

ART1-YFP constructs were made in pGREEN under the control of a 35S promoter. Constructs were infiltrated into 3-to 5-week-old *N. benthamiana*. After allowing 2 to 4 days for the protein to express, leaf discs were cut from tissue adjacent to the infiltration site and used for confocal microscopy. Confocal images were collected on a Leica TCS-SP5 confocal microscope (Leica Microsystems, Exton, PA) using a 63x water immersion objective. The YFP fusion protein was visualized by excitation with an argon ion laser (512 nm), and emitted light was collected between 525 nm and 575 nm. Chloroplasts were excited with the blue argon laser (488 nm), and emitted light was collected from 680 nm to 700 nm. The YFP and chloroplast fluorescence were collected on separate channels and superimposed with a bright field image collected simultaneously with the fluorescence images. Images were processed using Leica LAS-AF software (version 2.6.0).

### Yeast one hybrid experiments

A fragment of the *STAR1* promoter along with the five *ART1* alleles were cloned into pENTR D-T0PO (ThermoFisher, www.thermofisher.com). The *STAR1* promoter fragment (164 bp) was selected according to Yamaji et al (2009) who showed that this fragment contains an ART1 binding site. Clone identity was confirmed by Sanger sequencing, and the pENTR-STAR1p construct was sub-cloned into pLacZi-GW (Pruneda-Paz et al., 2014) by LR recombination, according to the manufacturer’s instructions (ThermoFisher). Bait strains were generated by homologous recombination of pLacZi-STAR1p into the yeast strain YM4271 according to manufacturer’s protocol (Clontech, www.clontech.com). Integration was verified by PCR. The prey vectors were generated by LR recombination of the pENTR-ART1 constructs into the yeast one-hybrid compatible plasmid pDEST22, which fuses the GAL4-Activation Domain to the N-terminus of the transcription factor. The bait strain was then transformed using the lithium acetate method (Gietz and Schiestl, 2007) with each of the five different pDEST22-ART1 constructs, and assayed for beta-galactosidase reporter gene activity using the liquid ONPG method described in the Clontech yeast protocol handbook (Clontech). Ten independent yeast colonies were screened for each allele. Student t-tests assuming unequal variance (and α=0.05) were performed to determine significant differences in DNA-binding affinity.

### RNA-Seq experiment and analysis

Plants were germinated in the dark in moist paper rolls for 3-4 days, then 40 uniform seedlings per genotype were grown in control nutrient solution for five days. Subsequently, 20 seedlings per genotype were transferred into a hydroponic solution with 80 μM of Al^3+^ activity for 4 hours before total root tissue was harvested for RNA extraction. Four independent biological replicates were performed, for a total of 32 samples (4 genotypes X 2 treatments X 4 biological replicates). Total RNA was isolated from root tissue using the QIAGEN RNeasy Mini Kit according to the manufacturer’s instructions (www.qiagen.com). Strand-specific RNA-Seq libraries (ssRNA-Seq) were constructed by Polar Genomics, as described by (Pombo et al., 2014) and (Zhong et al., 2011) (http://polargenomics.com/). Barcoded libraries were multiplexed by 16x for a total of 32 samples, and sequenced in two lanes of an Illumina Hiseq2000 using the single-end reads mode (Pombo et al. 2014). Raw RNA-Seq reads were processed to remove adaptor and low quality sequences using Trimmomatic (Bolger et al., 2014). RNA-Seq reads longer than 40 bp were kept. Reads were aligned to a ribosomal RNA database (Quast et al., 2013) using Bowtie (Langmead et al., 2009) and matches were discarded. The resulting, high-quality reads were aligned to the MSUv7.0 rice genome annotation (Kawahara et al., 2013) using HISAT (Kim et al., 2015). Following alignments, raw counts for each rice gene were derived and normalized to reads per Kilobase of exon model per million mapped reads (RPKM).

### Differential expression analysis

Differentially expressed (DE) genes were identified using the DESeq2 package (Love et al., 2014), https://bioconductor.org/packages/release/bioc/html/DESeq2.html). Independent filtering to remove low expression genes previous to DESeq2 analysis was performed by eliminating genes that across all samples their total sum counts is below the threshold estimated as: ***overall normalized mean counts / number of samples to be analyzed*** (Bourgon et al., 2010).

Differentially expressed genes (DEGs) were selected according to the following post-hoc criteria: **a)** two-fold or greater change in transcript abundance, up or down, measured in log2 fold-change ratio (log_2_FC ≥ 1, or ≤ -1); **b)** *p* value of ≤ 0.05, corrected for multiple testing using a false discovery rate (FDR) criteria (Benjamini and Hochberg, 1995); and **c)** minimum of eight normalized, log-transformed counts in at least one sample (Worley et al., 2016). Regularized log transformed counts (Love et al. 2014) estimated from DESeq2 using the true experimental design option (blind = TRUE; which accounts for different library sizes and stabilizes the variance among counts) were used to generate heat maps and for principal component analysis (PCA). For PCA, the variance across all samples for normalized, log-transformed counts was estimated using the “rowVars” function in R (https://cran.r-project.org/). Genes were ranked from the most to the least variance and the 500 genes with the highest variance across all samples were used to calculate principal components using the function “prcomp” in R (https://cran.r-project.org/), as described in Love et al. (2014). Heat maps were generated in R using the package “gplots” (http://cran.stat.nus.edu.sg/web/packages/gplots/gplots.pdf) and the function ‘heat map.2’. Principal component analysis (PCA) was carried out using the regularized-logarithm transformed counts from DESeq2. The control samples NIL AZU_IR6412.1_ rep 3, IR64 rep 4 and NIL-IR64_AZU12.1_ rep 1, as well as Al-treated samples NIL-AZU_IR6412.1_ rep 3, IR64 rep 3, and NIL-IR64_AZU12.1_ rep 1 did not group near their respective genotype/treatment samples in the PCA analysis, and therefore were removed from the PCA shown in Suppl. Figure 3. Nevertheless, all biological replicates were used for differential gene expression analysis. GO enrichment analysis was performed using Plant MetGenMAP (http://bioinfo.bti.cornell.edu/cgi-bin/MetGenMAP/home.cgi). We applied an FDR-corrected *p* value cut-off of 0.01.

## Supplemental Material

**Supplemental Figure S1:** Fine-mapping of the Al resistance QTL *Alt12.1*.

**Supplemental Figure S2:** Alignment of the QTL *Alt12.1* fine-mapped region (Nipponbare reference genome) to IR64 (PacBio assembly).

**Supplemental Figure S3:** Principal component analysis (PCA) of gene expression profiles. Supplemental Figure S4: Differential expression of LOC_Os12g07340, located within the *Alt12.1* fine-mapped region, based on RNA-seq.

**Supplemental Figure S5:** GO analysis of down-regulated genes in response to Al in Azucena (left) and IR64 (right) backgrounds.

**Supplemental Figure S6:** The presence of the *ART1* allele from Azucena may not affect the same set of genes in *indica* and *japonica* backgrounds.

**Supplemental Figure S7:** *OsFRDL4* expression is affected by the different ART1 alleles in Azucena and IR64 genetic backgrounds.

**Supplemental Figure S8:** Occurrence of ART1 predicted binding sites in putative regulatory regions of rice genes.

**Supplemental Methods S1:** Developing of *Alt12.1* reciprocal near-isogenic lines (NILs). Supplemental Methods S2: Fine mapping of *Alt12.1*.

**Supplemental Table S1:** Genomic composition and Al resistance phenotype of the reciprocal NILs.

**Supplemental Table S2:** Molecular markers used during fine-mapping analysis.

**Supplemental Table S3:** Statistics for the RNA-seq read alignments.

**Supplemental Table S4:** SNPs genotyped for RNA-seq sample ID verification across 14 loci in Azucena, IR64, and the NILs AZU_[IR6412.1]_ and IR64_[AZU12.1]_.

## Accession numbers

Sequences of the *ART1* alleles for Azucena, IR64, Nipponbare and Kasalath were submitted to GenBank (accession numbers MF093140 – MF093143).

Illumina RNA-Seq reads were submitted to Gene Expression Omnibus (GEO; www.ncbi.nlm.nih.gov/geo/info/seq.html) at the National Center for Biotechnology Information (NCBI; www.ncbi.nlm.nih.gov/) under the accession number GSE89494.

The *de novo* IR64 genome assembly is available at http://schatzlab.cshl.edu/data/ir64/.

## Large Datasets

**Supplemental File S1.** Genotypic data for Azucena, IR64, and the NILs AZU_[IR6412.1]_ and IR64_[AZU12.1]_ using 2410 polymorphic SNPs from the Infinium 6K Illumina platform (Thomson et al. 2016). The position and length of each reciprocal introgression on chromosome 12 is indicated in each NIL.

**Supplemental File S2.** Differentially-regulated genes in the RNA-seq study.

**Supplemental File S3.** GO enrichment analysis of down-regulated genes.

**Supplemental File S4.** Comparison of the fold-change of differentially-regulated genes in each genetic background.

## Acknowledgments

We acknowledge the following people without whom the completion of this work would not have been possible: We are grateful to Sandy Harrington, Gen Onishi, Joseph Gage, James Jones-Rounds, Anjali T. Merchant, Yuxin Xi, Francisco Agosto-Perez, Eric Craft and Alison Coluccio for technical and management support; and to Karl Kremling and Christine Diepenbrock for critically reviewing a draft of the manuscript.

## Author contributions

JDA, LGM, AF, TOJ, ARR, MAP and NS performed experiments; LGM, JDA, AF, QM and ZF analyzed data; LGM, JDA, and SRM wrote the paper; SRM, AF, JDA, LGM, and LVK conceived the project.

## Notes

**Funding information:** this work was supported by a grant from USDA-NIFA, award # 201367013-21379; NSF-PGRP award #1026555; a PhD fellowship for JDA from the Monsanto Beachell-Bourlag International Scholars Program.

## Literature Cited

Arbelaez JD, Moreno LT, Singh N, Tung CW, Maron LG, Ospina Y, Martinez CP, Grenier C, Lorieux M, McCouch S (2015) Development and GBS-genotyping of introgression lines (ILs) using two wild species of rice, and in a common recurrent parent, cv. Curinga. Mol Breed 35: 81

Arenhart RA, Bai Y, Valter de Oliveira LF, Bucker Neto L, Schunemann M, Maraschin FdS, Mariath J, Silverio A, Sachetto-Martins G, Margis R, Wang Z-Y, Margis-Pinheiro M (2014) New Insights into Aluminum Tolerance in Rice: The ASR5 Protein Binds the *STAR1* Promoter and other Aluminum-Responsive Genes. Molecular Plant 7: 709–721

Benjamini Y, Hochberg Y (1995) Controlling the False Discovery Rate - A Practical and Powerful Approach to Multiple Testing. Journal of the Royal Statistical Society Series B-Methodological 57: 289–300

Bolger AM, Lohse M, Usadel B (2014) Trimmomatic: a flexible trimmer for Illumina sequence data. Bioinformatics (Oxford, England) 30: 2114–2120

Bourgon R, Gentleman R, Huber W (2010) Independent filtering increases detection power for high-throughput experiments. Proceedings of the National Academy of Sciences of the United States of America 107: 9546–9551

Chandran D, Sharopova N, Ivashuta S, Gantt JS, VandenBosch KA, Samac DA (2008) Transcriptome profiling identified novel genes associated with aluminum toxicity, resistance and tolerance in Medicago truncatula. Planta 228: 151–166

Clark RT, Famoso AN, Zhao K, Shaff JE, Craft EJ, Bustamante CD, McCouch SR, Aneshansley DJ, Kochian LV (2013) High-throughput two-dimensional root system phenotyping platform facilitates genetic analysis of root growth and development. Plant, Cell & Environment 36: 454–466

Famoso AN, Clark RT, Shaff JE, Craft E, McCouch SR, Kochian LV (2010) Development of a novel aluminum tolerance phenotyping platform used for comparisons of cereal aluminum tolerance and investigations into rice aluminum tolerance mechanisms. Plant Physiology 153: 1678–1691

Famoso AN, Zhao K, Clark RT, Tung C-W, Wright MH, Bustamante C, Kochian LV, McCouch SR (2011) Genetic Architecture of Aluminum Tolerance in Rice (*Oryza sativa*) Determined through Genome-Wide Association Analysis and QTL Mapping. PLoS Genet 7: e1002221

Foy CD (1988) Plant adaptation to acid, aluminum-toxic soils. Commun Soil Sci Plant Anal 19: 959–987

Franco-Zorrilla JM, López-Vidriero I, Carrasco JL, Godoy M, Vera P, Solano R (2014) DNA-binding specificities of plant transcription factors and their potential to define target genes. Proceedings of the National Academy of Sciences 111: 2367–2372

Garris AJ, Tai TH, Coburn J, Kresovich S, McCouch S (2005) Genetic structure and diversity in *Oryza sativa* L. Genetics 169: 1631–1638

Gietz RD, Schiestl RH (2007) High-efficiency yeast transformation using the LiAc/SS carrier DNA/PEG method. Nat. Protocols 2: 31–34

Huang C-F, Yamaji N, Chen Z, Ma JF (2012) A tonoplast-localized half-size ABC transporter is required for internal detoxification of aluminum in rice. The Plant Journal 69: 857–867

Huang CF, Yamaji N, Mitani N, Yano M, Nagamura Y, Ma JF (2009) A Bacterial-Type ABC Transporter Is Involved in Aluminum Tolerance in Rice. The Plant Cell 21: 655–667

Imai I, Kimball J, Conway B, Yeater K, McCouch S, McClung A (2013) Validation of yield-enhancing quantitative trait loci from a low-yielding wild ancestor of rice. Molecular Breeding 32: 101–120

Iuchi S, Koyama H, Iuchi A, Kobayashi Y, Kitabayashi S, Kobayashi Y, Ikka T, Hirayama T, Shinozaki K, Kobayashi M (2007) Zinc finger protein STOP1 is critical for proton tolerance in Arabidopsis and coregulates a key gene in aluminum tolerance. Proceedings of the National Academy of Sciences 104: 9900–9905

Kawahara Y, de la Bastide M, Hamilton JP, Kanamori H, McCombie WR, Ouyang S, Schwartz DC, Tanaka T, Wu J, Zhou S, Childs KL, Davidson RM, Lin H, Quesada-Ocampo L, Vaillancourt B, Sakai H, Lee SS, Kim J, Numa H, Itoh T, Buell CR, Matsumoto T, Cingolani P, Platts A, Wang LL, Coon M, Nguyen T, Wang L, Land SJ, Lu X, Ruden DM, Eichten SR, Foerster JM, Leon Nd, Kai Y, Yeh CT, Liu S, Jeddeloh JA, Schnable PS, Kaeppler SM, Springer NM, Goff SA, Ricke D, Lan TH, Presting G, Wang R, Dunn M, Glazebrook J, Sessions A, Oeller P, Varma H, Hadley D, Hutchison D, Martin C, Katagiri F, Lange BM, Moughamer T, Xia Y, Budworth P, Zhong J, Miguel T, Paszkowski U, Zhang S, Colbert M, Sun WL, Chen L, Cooper B, Park S, Wood TC, Mao L, Quail P, Huang X, Wei X, Sang T, Zhao Q, Feng Q, Zhao Y, Li C, Zhu C, Lu T, Zhang Z, Li M, Fan D, Guo Y, Wang A, Wang L, Deng L, Li W, Lu Y, Weng Q, Liu K, Huang T, Zhou T, Jing Y, Li W, Lin Z, Buckler ES, Qian Q, Zhang QF, Li J, Han B, Huang X, Zhao Y, Wei X, Li C, Wang A, Zhao Q, Li W, Guo Y, Deng L, Zhu C, Fan D, Lu Y, Weng Q, Liu K, Zhou T, Jing Y, Si L, Dong G, Huang T, Lu T, Feng Q, Qian Q, Li J, Han B, Li H, Durbin R, Li H, Handsaker B, Wysoker A, Fennell T, Ruan J, Homer N, Marth G, Abecasis G, Lu T, Lu G, Fan D, Zhu C, Li W, Zhao Q, Feng Q, Zhao Y, Guo Y, Li W, Huang X, Han B, McMullen MD, Kresovich S, Villeda HS, Bradbury P, Li H, Sun Q, Flint-Garcia S, Thornsberry J, Acharya C, Bottoms C, Brown P, Browne C, Eller M, Guill K, Harjes C, Kroon D, Lepak N, Mitchell SE, Peterson B, Pressoir G, Romero S, Rosas MO, Salvo S, Yates H, Hanson M, Jones E, Smith S, Glaubitz JC, Goodman M, Ware D, Murray MG, Thompson WF, Ohmido N, Kijima K, Akiyama Y, Jong JHd, Fukui K, Oono K, Sugiura M, Ossowski S, Schneeberger K, Lucas-Lledó JI, Warthmann N, Clark RM, Shaw RG, Weigel D, Lynch M, Ouyang S, Zhu W, Hamilton J, Lin H, Campbell M, Childs K, Thibaud-Nissen F, Malek RL, Lee Y, Zheng L, Orvis J, Haas B, Wortman J, Buell CR, Sakai H, Itoh T, Spannagl M, Noubibou O, Haase D, Yang L, Gundlach H, Hindemitt T, Klee K, Haberer G, Schoof H, Mayer KFX, Stein LD, Mungall C, Shu S, Caudy M, Mangone M, Day A, Nickerson E, Stajich JE, Harris TW, Arva A, Lewis S, Tanaka T, Antonio BA, Kikuchi S, Matsumoto T, Nagamura Y, Numa H, Sakai H, Wu J, Itoh T, Sasaki T, Aono R, Fujii Y, Habara T, Harada E, Kanno M, Kawahara Y, Kawashima H, Kubooka H, Matsuya A, Nakaoka H, Saichi N, Sanbonmatsu R, Sato Y, Shinso Y, Suzuki M, Takeda JI, Tanino M, Todokoro F, Yamaguchi K, Yamamoto N, Teague B, Waterman MS, Goldstein S, Potamousis K, Zhou S, Reslewic S, Sarkar D, Valouev A, Churas C, Kidd JM, Kohn S, Runnheim R, Lamers C, Forrest D, Newton MA, Eichler EE, Kent-First M, Surti U, Livny M, Schwartz DC, Valouev A, Li L, Liu Y, Schwartz DC, Yang Y, Zhang Y, Waterman MS, Xu X, Liu X, Ge S, Jensen JD, Hu F, Li X, Dong Y, Gutenkunst RN, Fang L, Huang L, Li J, He W, Zhang G, Zheng X, Zhang F, Li Y, Yu C, Kristiansen K, Zhang X, Wang J, Wright M, McCouch S, Nielsen R, Wang J, Wang W, Yamamoto T, Nagasaki H, Yonemaru JI, Ebana K, Nakajima M, Shibaya T, Yano M, Yang CC, Kawahara Y, Mizuno H, Wu J, Matsumoto T, Itoh T, Zhang Z, Schwartz S, Wagner L, Miller W, Zhang G, Guo G, Hu X, Zhang Y, Li Q, Li R, Zhuang R, Lu Z, He Z, Fang X, Chen L, Tian W, Tao Y, Kristiansen K, Zhang X, Li S, Yang H, Wang J, Wang J, Zhou S, Bechner MC, Place M, Churas CP, Pape L, Leong SA, Runnheim R, Forrest DK, Goldstein S, Livny M, Schwartz DC (2013) Improvement of the Oryza sativa Nipponbare reference genome using next generation sequence and optical map data. Rice 6: 4–4

Kim D, Langmead B, Salzberg SL (2015) HISAT: a fast spliced aligner with low memory requirements. Nature Methods 12: 357–360

Kochian LV, Piñeros MA, Liu J, Magalhaes JV (2015) Plant Adaptation to Acid Soils: The Molecular Basis for Crop Aluminum Resistance. Annual Review of Plant Biology 66: 571–598

Langmead B, Trapnell C, Pop M, Salzberg SL (2009) Ultrafast and memory-efficient alignment of short DNA sequences to the human genome. Genome biology 10: R25–R25

Li J-Y, Liu J, Dong D, Jia X, McCouch SR, Kochian LV (2014) Natural variation underlies alterations in Nramp aluminum transporter (NRAT1) expression and function that play a key role in rice aluminum tolerance. Proceedings of the National Academy of Sciences 111: 6503–6508

Love MI, Huber W, Anders S (2014) Moderated estimation of fold change and dispersion for RNA-seq data with DESeq2. Genome Biology 15: 550–550

Ma JF, Shen R, Zhao Z, Wissuwa M, Takeuchi Y, Ebitani T, Yano M (2002) Response of Rice to Al Stress and Identification of Quantitative Trait Loci for Al Tolerance. Plant and Cell Physiology 43: 652–659

Maron LG, Kirst M, Mao C, Milner MJ, Menossi M, Kochian LV (2008) Transcriptional profiling of aluminum toxicity and tolerance responses in maize roots. New Phytologist 179: 116–128

Maron LG, Piñeros MA, Kochian LV, McCouch SR (2016) Redefining ‘stress resistance genes’, and why it matters. Journal of Experimental Botany 67: 5588–5591

Melo JO, Lana UGP, Piñeros MA, Alves VMC, Guimarães CT, Liu J, Zheng Y, Zhong S, Fei Z, Maron LG, Schaffert RE, Kochian LV, Magalhaes JV (2013) Incomplete transfer of accessory loci influencing SbMATE expression underlies genetic background effects for aluminum tolerance in sorghum. The Plant Journal 73: 276–288

Mickelbart MV, Hasegawa PM, Bailey-Serres J (2015) Genetic mechanisms of abiotic stress tolerance that translate to crop yield stability. Nat Rev Genet 16: 237–251

Mora-Macías J, Ojeda-Rivera JO, Gutiérrez-Alanís D, Yong-Villalobos L, Oropeza-Aburto A, Raya-González J, Jiménez-Domínguez G, Chávez-Calvillo G, Rellán-Álvarez R, Herrera-Estrella L (2017) Malate-dependent Fe accumulation is a critical checkpoint in the root developmental response to low phosphate. Proceedings of the National Academy of Sciences 114: E3563–E3572

Pombo MA, Zheng Y, Fernandez-Pozo N, Dunham DM, Fei Z, Martin GB (2014) Transcriptomic analysis reveals tomato genes whose expression is induced specifically during effector-triggered immunity and identifies the Epk1 protein kinase which is required for the host response to three bacterial effector proteins. Genome Biology 15

Pruneda-Paz Jose L, Breton G, Nagel Dawn H, Kang SE, Bonaldi K, Doherty Colleen J, Ravelo S, Galli M, Ecker Joseph R, Kay Steve A (2014) A Genome-Scale Resource for the Functional Characterization of Arabidopsis Transcription Factors. Cell Reports 8: 622–632

Quast C, Pruesse E, Yilmaz P, Gerken J, Schweer T, Yarza P, Peplies J, Glöckner FO (2013) The SILVA ribosomal RNA gene database project: improved data processing and web-based tools. Nucleic acids research 41: D590–596

Rombauts S, Florquin K, Lescot M, Marchal K, Rouzé P, Van de Peer Y (2003) Computational Approaches to Identify Promoters and cis-Regulatory Elements in Plant Genomes. Plant Physiology 132: 1162–1176

Spindel J, Wright M, Chen C, Cobb J, Gage J, Harrington S, Lorieux M, Ahmadi N, McCouch S (2013) Bridging the genotyping gap: using genotyping by sequencing (GBS) to add high-density SNP markers and new value to traditional bi-parental mapping and breeding populations. Theoretical and Applied Genetics 126: 2699–2716

Tsutsui T, Yamaji N, Feng Ma J (2011) Identification of a Cis-Acting Element of ART1, a C2H2-Type Zinc-Finger Transcription Factor for Aluminum Tolerance in Rice. Plant Physiology 156: 925–931

Tsutsui T, Yamaji N, Huang CF, Motoyama R, Nagamura Y, Ma JF (2012) Comparative Genome-Wide Transcriptional Analysis of Al-Responsive Genes Reveals Novel Al Tolerance Mechanisms in Rice. PLoS ONE 7: e48197

Vaahtera L, Brosché M (2011) More than the sum of its parts – How to achieve a specific transcriptional response to abiotic stress. Plant Science 180: 421–430

Vandepoele K, Quimbaya M, Casneuf T, De Veylder L, Van de Peer Y (2009) Unraveling Transcriptional Control in Arabidopsis Using cis-Regulatory Elements and Coexpression Networks. Plant Physiology 150: 535–546

Vázquez-Lobo A, Roujol D, Zuñiga-Sánchez E, Albenne C, Piñero D, Buen AGd, Jamet E (2012) The highly conserved spermatophyte cell wall DUF642 protein family: Phylogeny and first evidence of interaction with cell wall polysaccharides in vitro. Molecular Phylogenetics and Evolution 63: 510–520

von Uexküll HR, Mutert E (1995) Global Extent, Development and Economic-Impact of Acid Soils. Plant and Soil 171: 1–15

Worley JN, Pombo MA, Zheng Y, Dunham DM, Myers CR, Fei Z, Martin GB (2016) A novel method of transcriptome interpretation reveals a quantitative suppressive effect on tomato immune signaling by two domains in a single pathogen effector protein. BMC Genomics 17: 229–229

Xia J, Yamaji N, Kasai T, Ma JF (2010) Plasma membrane-localized transporter for aluminum in rice. Proceedings of the National Academy of Sciences of the United States of America 107: 18381–18385

Xia J, Yamaji N, Ma JF (2013) A plasma membrane-localized small peptide is involved in rice aluminum tolerance. The Plant Journal 76: 345–355

Yamaji N, Huang CF, Nagao S, Yano M, Sato Y, Nagamura Y, Ma JF (2009) A Zinc Finger Transcription Factor ART1 Regulates Multiple Genes Implicated in Aluminum Tolerance in Rice. The Plant Cell 21: 3339–3349

Yokosho K, Yamaji N, Kashino-Fujii M, Ma JF (2016) Retrotransposon-mediated aluminum tolerance through enhanced expression of the citrate transporter OsFRDL4. Plant Physiology 172: 2327–2336

Yokosho K, Yamaji N, Ma JF (2011) An Al-inducible MATE gene is involved in external detoxification of Al in rice. The Plant Journal 68: 1061–1069

Yokosho K, Yamaji N, Ma JF (2014) Global Transcriptome Analysis of Al-Induced Genes in an Al-Accumulating Species, Common Buckwheat (Fagopyrum esculentum Moench). Plant and Cell Physiology 55: 2077–2091

Zhao K, Tung C-W, Eizenga GC, Wright MH, Ali ML, Price AH, Norton GJ, Islam MR, Reynolds A, Mezey J, McClung AM, Bustamante CD, McCouch SR (2011) Genome-wide association mapping reveals a rich genetic architecture of complex traits in *Oryza sativa*. Nat Commun 2: 467

Zhong S, Joung J-G, Zheng Y, Chen Y-r, Liu B, Shao Y, Xiang JZ, Fei Z, Giovannoni JJ (2011) High-Throughput Illumina Strand-Specific RNA Sequencing Library Preparation. Cold Spring Harbor Protocols 2011: pdb.prot5652

Zúñiga-Sánchez E, Soriano D, Martínez-Barajas E, Orozco-Segovia A, Gamboa-deBuen A (2014) BIIDXI, the At4g32460 DUF642 gene, is involved in pectin methyl esterase regulation during Arabidopsis thaliana seed germination and plant development. BMC Plant Biology 14: 338

